# RNA Pol I activity is required for meiotic chromatin organization and the H3K4me3 gradient essential for oogenesis, independent of ribosome synthesis

**DOI:** 10.1101/2025.05.07.652530

**Authors:** Raquel Mejia-Trujillo, Qiuxia Zhao, Francesca Abraham, Ayesha Rahman, Elif Sarinay Cenik

## Abstract

Oogenesis requires extensive and dynamic chromatin remodeling that primes gene promoters for later transcriptional activation during embryonic development. Here, we uncover a pivotal, non-canonical role for RNA Polymerase I (Pol I) in driving these chromatin state transitions during *Caenorhabditis elegans* oogenesis. Using the auxin-inducible degron system to selectively deplete either Pol I catalytic subunits or ribosome assembly factors, we disentangle the consequences of impaired nucleolar integrity from reductions in ribosome biogenesis. Strikingly, although disrupting ribosome assembly caused minimal effects on oocyte production, loss of Pol I activity led to widespread changes in chromatin accessibility, a dampening of the distal-proximal H3K4me3 gradient required for oogenesis, reduced synapsis, and elevated ATM/ATR phosphorylation, resulting in fewer but significantly larger oocytes. Despite their promoters becoming more accessible, oogenesis genes did not show large changes in steady-state mRNA, consistent with transcriptional repression prior to fertilization. Instead, Pol I depletion prematurely remodeled oogenic chromatin, through a misdirection of H3K4me3 deposition towards promoters normally primed for zygotic genome activation. These findings reveal an epigenetic gating function for nucleolar integrity in oocyte maturation: Pol I preserves three-dimensional chromatin organization and maintains proper spatiotemporal regulation of histone modifications, independent of ribosome production. Given the evolutionary conservation of nucleolar dynamics and histone modifications during gametogenesis, our work suggests that nucleolar stress, whether from environmental factors, aging, or genetic disorders, could broadly compromise fertility by disrupting oogenic chromatin priming.

**GRAPHICAL ABSTRACT:** 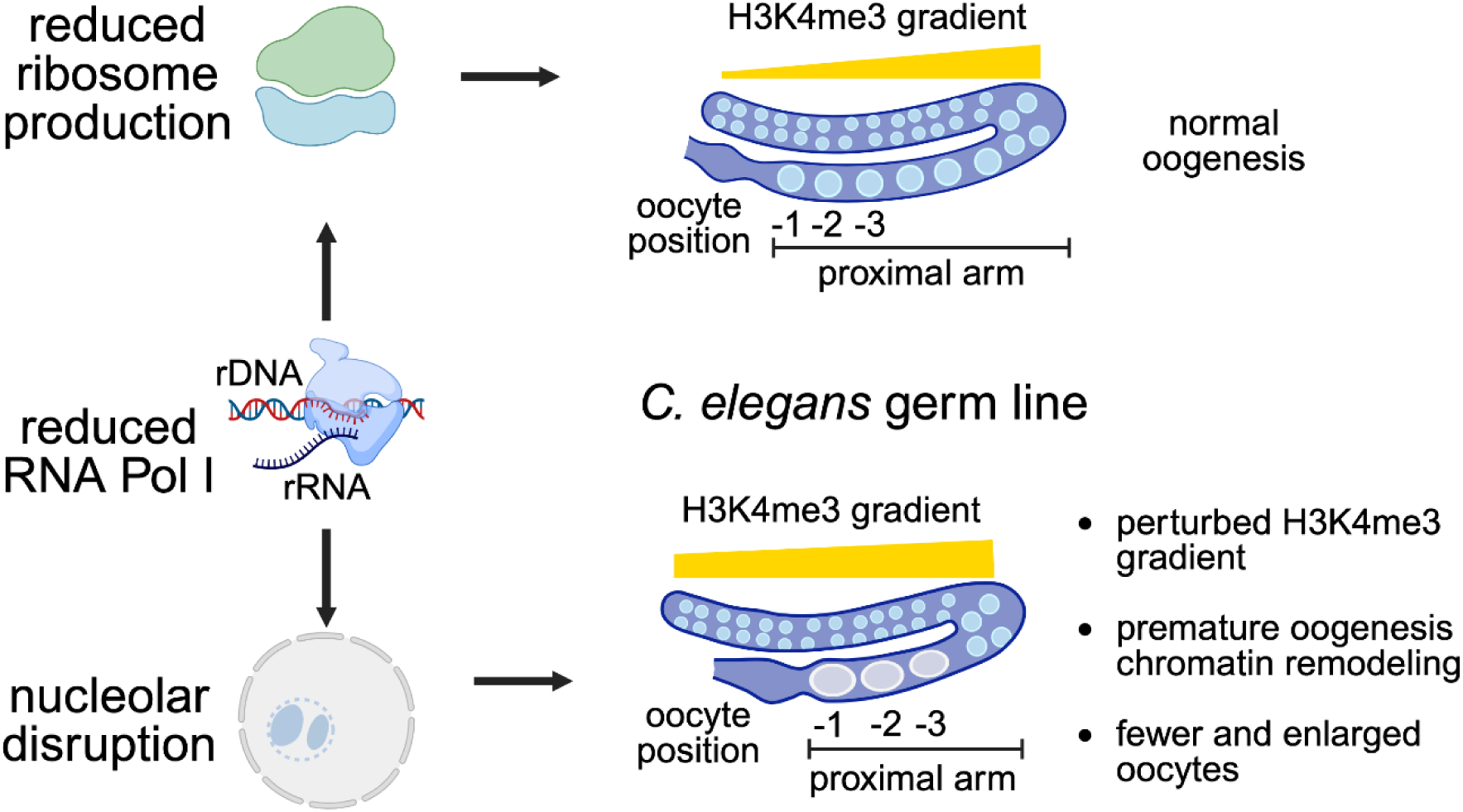

## INTRODUCTION

Chromosome condensation and chromatin modifications during meiosis are essential to ensure accurate genetic transmission in sexually reproducing species. In the hermaphroditic germ line of *Caenorhabditis elegans*, female gamete production proceeds through four distinct zones: mitotic proliferation, transition, meiosis I, and oocyte maturation. Upon completion of oocyte maturation, the most proximal oocyte is fertilized by the adjacent spermatheca, initiating zygotic development.

Oogenesis involves a complex progression of germ cell maturation, yet net transcriptional activity is strikingly low in late-stage oocytes. Instead, many oogenesis genes remain largely quiescent before fertilization, with regulation relying primarily on extensive chromatin remodeling. This epigenetic priming prepares the promoters of key developmental genes to be robustly activated after fertilization or during early embryogenesis ^1–3^.

In *C. elegans*, as in other species, germ cells undergo profound chromatin reorganization during meiotic progression to form specialized meiotic chromatin structures, establish parental imprinting marks, and position active marks near the promoters of genes that will be engaged during zygotic genome activation (ZGA) ^4^. Specifically, H3K4me3 marks shift from gene bodies to promoters as undifferentiated mitotic germ cells progress through meiosis I ^4^. This promoter-specific priming of ZGA genes with H3K4me3 is evolutionarily conserved across species, including *Xenopus* ^5^, zebrafish ^6,7^, mice ^8,9^, and humans ^10^ and is critical for normal germ line development. For instance, *C. elegans* germ lines lacking H3K4 methyltransferase activity exhibit reduced oocyte production, aberrant nuclear morphology, and aneuploid gametes ^11,12^, while deficient H3K4 demethylase activity leads to abnormal chromatin condensation, diminished oocyte production, and failed entry into meiosis ^13^.

In parallel with the spatiotemporal regulation of H3K4 methylation, maturing germ cells also temporally regulate nucleolar activity during oogenesis. The nucleolus, responsible for rRNA synthesis and ribosome assembly ^14^, remains intact throughout meiosis but temporarily disassembles at the onset of oocyte maturation before reappearing during ZGA ^15^. Similar nucleolar dynamics have been documented across hermaphrodite and dioecious eukaryotes, including *C. elegans*, yeast, pig, and mouse ^15–20^. The presence of the nucleolus throughout oogenesis is unsurprising ^21^, given that maternal ribosome production is sufficient to drive embryogenesis in *Caenorhabditis elegans* ^22^; however, the nucleolus may have additional roles beyond ribosome production during oogenesis. For example, in *Arabidopsis*, rDNA loci are embedded within the nucleolus to prevent alterations of rDNA copy numbers and genomic instability during meiotic homologous recombination ^23^. In mouse oocytes, maternal nucleolar components are essential for maintaining satellite repeat sequences immediately after fertilization ^24^. Lastly, the nucleolar microenvironment in *C. elegans* can sequester transcriptional silencing factors, such as PIE-1, away from the genome to promote transcription during late oogenesis ^25^.

Despite these hints of broader functions for the nucleolus, dissecting its direct contributions to oogenesis has been challenging because rRNA synthesis, ribosome assembly, and the maintenance of a spatially distinct nuclear subcompartment are all interconnected. For example, RNA Polymerase I (Pol I) transcription of rDNA loci is accompanied by co-transcriptional processing of pre-rRNA, followed by co-assembly with ribosomal proteins into ribosome subunits ^26^. The nucleolar microenvironment is similarly co-dependent on these other functions, as its liquid-liquid phase-separated properties make its spherical morphology reliant on active rDNA transcription and subsequent processing and assembly ^27,28^. Thus, similar to how separation-of-function mutations in proteins are used to differentiate between distinct protein functions ^29–31^, there is a demand for genetic tools that can isolate interconnected functions of the same subcellular structure. Here, we utilized separation-of-function *C. elegans* strains to identify potential nucleolar functions that are independent from ribosome production during oogenesis.

Using the auxin-inducible degron (AID) system, we targeted either two catalytic subunits specific to RNA Pol I (orthologous to human *POL1RA* and *POL1RB*) or two ribosome assembly factors essential for 60S and 40S assembly (orthologous to human *GRWD1* and *TSR2,* respectively). This approach isolates the impact of nucleolar disruption on female germ cell maturation from perturbed downstream ribosome assembly. Our results uncover a non-ribosomal function of RNA Pol I in regulating oocyte production, meiotic chromosome morphology and chromatin landscape, and the maintenance of an essential H3K4 methylation gradient. Germ cells fail to regulate chromatin remodeling during the transition from mitotic proliferation to meiotic prophase I following reduced RNA Pol I activity, but not after a decrease in ribosome assembly. ATAC-seq analyses further reveal that chromatin accessibility increases near oogenesis promoters, oogenesis-promoting EFL-1-binding sites, and promoters subject to canonical H3K4me3 remodeling during oogenesis. Nucleolar disruption induced upon RNA Pol I depletion also dampened the distal-proximal H3K4me3 gradient through premature H3K4me3 deposition and resulted in increased ATM/ATR phosphorylation in the pachytene region of the germ line. Collectively, our data suggests that nucleolar integrity, maintained by RNA Pol I activity, is involved in the coordination of chromatin remodeling during oogenesis, ensuring the production of healthy, properly sized oocytes.

## RESULTS

### Persistence and dynamics of nucleolar structure during *Caenorhabditis elegans* germ line development and oogenesis

We first examined nucleolar morphology in a wild-type background to establish a baseline representation of its structure during germ line development, from the mitotic through meiotic zones, and into oocyte maturation. To accomplish this, we fluorescently tagged two genes: *nucl-1*, the *Caenorhabditis elegans* ortholog of human *NCL*, which facilitates rRNA transcription and processing ^32–34^, and *dao-5*, the ortholog of human *NOLC1*, required for nucleologenesis and rDNA transcription ^15,35^. We also used a fluorescent SYP-3 reporter, a key component of the synaptonemal complex ^36,37^, to distinguish between different germ line zones. Cells distal to synaptonemal complex formation were classified as mitotic, cells displaying a synaptonemal complex were assigned to the transition or pachytene zone, and cells with visible plasma membrane under transmitted photomultiplier tubes (T-PMT) were defined as being in the oocyte maturation zone.

In a mature germ line, NUCL-1 retains a spherical nucleolar localization pattern in the mitotic, meiotic, and most of the oocyte maturation zone (**Figure S1A**). As the oocytes matured, the nucleolus formed intermediate droplet-like structures in the “-2” oocyte (the second-most proximal oocyte), followed by complete dissolution in the “-1” oocyte (the most proximal one) (**Figure S1B**). These results suggest that nucleolar integrity is largely maintained from the distal mitotic region through meiosis but is disrupted just before fertilization. This observation agrees with previous reports in *C. elegans* germ lines ^15,38,39^ and is also conserved in mammalian oocytes ^16,18–20^.

### Disruption of RNA Pol I activity compromises nucleolar structure throughout the germ line

Having established that a spherical nucleolus persists through most of germ line development in mature hermaphrodites, we next asked how reducing rDNA transcription would affect its structure. Nucleolar integrity depends on robust RNA polymerase I (Pol I) activity, as the formation of nucleolar subcompartments and phase separation properties are largely driven by elevated concentrations of rRNA, along with its bound nucleolar factors ^27,28,40^. Although prior work in the *C. elegans* germ line utilized Actinomycin D to inhibit rDNA transcription, no visible disturbance of nucleolar structure was detected, likely because transcription was not suppressed below a critical threshold associated with nucleolar segregation ^41,42^. Thus, to clearly disrupt nucleolar structure, we used the auxin-inducible degron (AID) system to substantially deplete two endogenously tagged subunits of RNA Pol I: RPOA-1, the largest subunit of RNA Pol I (orthologous to human *POLR1A*), and RPOA-2, a catalytic subunit specific to RNA Pol I (orthologous to human *POLR1B*) ^38,43^.

Animals expressing degron::GFP::RPOA-2 and RPOA-1::degron::GFP were treated with 1 mM IAA (auxin) at the L4 stage for 18 hours, after which gonads from young adults were dissected and examined by fluorescent microscopy. A significant decrease in mean GFP intensity confirmed a substantial depletion of both RPOA-1 and RPOA-2 in germ cells (**Figure 1A-B** and **1E**). To confirm that this depletion impaired rDNA transcription, we performed RNA-seq on gonadal samples using total RNA and quantified the abundance of reads mapping to two loci uniquely found in nascent pre-rRNA. We observed a significant decrease in reads mapping to the internally transcribed spacers (ITS1 and ITS2), indicating a considerable decrease in RNA Pol I activity upon depletion of RPOA-2 (**Figure 1F** and **S1C**, one-tailed Welch two-sample *t*-test, 4.6-fold average decrease for ITS1: *p* = 0.021, 1.6-fold average reduction for ITS2: *p* = 0.0092). RT-qPCR also supports that the depletion of RPOA-2 significantly reduces pre-rRNA synthesis **(Figure S1D**, left-tailed paired Student’s *t-*test, ITS1: *p* = 0.0048, ITS2: *p* = 0.0015).

**Figure 1.**
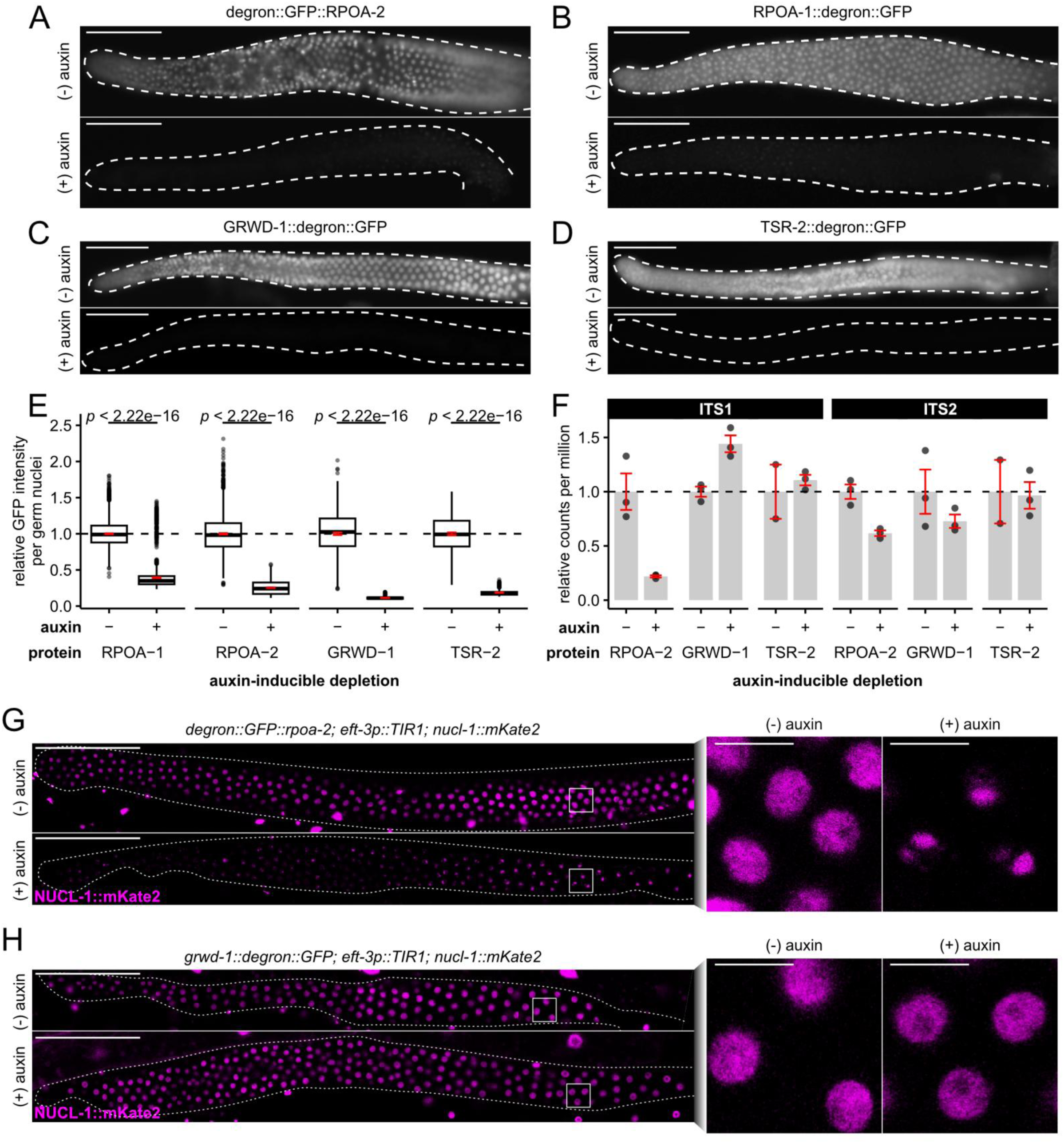
Auxin-inducible depletion of RNA Pol I subunits and ribosome assembly factors reveal that nucleolar structure is dependent on pre-rRNA synthesis but not downstream ribosome maturation. See also Figure S1. (**A-D**) Germline expression of fluorescently tagged RPOA-2 (**A**), RPOA-1 (**B**), GRWD-1 (**C**), and TSR-2 (**D**) proteins after control or auxin treatment. Dissected germ lines are outlined in white. Each strain’s control/auxin images were taken with identical exposure settings. Scale bar = 50 μm. (**E**) Distribution of germ nuclei GFP intensity after auxin-inducible depletion of GFP-tagged RPOA-1/RPOA-2/GRWD-1/TSR-2. Statistical comparisons performed using left-tailed Welch’s two-sample *t*-tests. (**F**) Comparisons of counts per million (CPM) reads across a simplified single copy of the ITS1 and ITS2 loci based on gonadal RNA-seq (total RNA) after auxin-inducible depletion of RPOA-2/GRWD-1/TSR-2. Points represent biological replicates (3 per condition, each consisting of 20 gonads); bars and error bars represent the mean and SEM, respectively. (**G-H**) Germline nucleolar morphology (endogenous NUCL-1::mKate2) after auxin-inducible depletion of RPOA-2 (**G**) or GRWD-1 (**H**). Germ lines are outlined in white; zoomed-in nucleoli are shown in the right column. Scale bars = 50 μm (whole gonad) and 5 μm (zoomed nucleoli).

We next investigated whether these transcriptional reductions could sufficiently disrupt spherical nucleolar structure, identifiable by the formation of nucleolar caps ^42^. In RPOA-2-depleted gonads, the nucleolar marker NUCL-1::mKate2, showed droplet and cap-like structures throughout the germ line, in contrast to the more uniform spherical nucleoli in control animals (**Figure 1G**). This phenotype suggests that the threshold of rDNA transcription required for nucleolar integrity was not met upon RPOA-2 depletion, leading to a clear disruption of nucleolar morphology. We validated that auxin treatment alone did not induce any unintended morphological changes to gonadal NUCL-1::mKate2 in the absence of the AID system (**Figure S1F**).

Although RPOA-2 depletion disrupts nucleolar structure by reducing rRNA synthesis, it also suspends ribosome production, which might impact the translational capacity of developing germ cells ^43^. Despite the generally low transcriptional activity of germ cells, proper ribosome production is essential for translational regulation of gene expression and maternal ribosome loading to ensure embryonic viability ^22,44–46^. Importantly, our prior work indicates that both the depletion of RPOA-2 or ribosome assembly factors result in a comparable decrease in overall ribosome production, as evidenced by developmental arrest on auxin ^43^. Thus, to separate the effects of nucleolar disruption from potential consequences of ribosome biogenesis disruption, we conducted identical experiments using endogenous fluorescently-labeled AID strains for GRWD-1, a chaperone that guides RPL-3 assembly into the 60S ribosomal subunit ^43,47^, and TSR-2, a chaperone that incorporates RPS-26 into the 40S subunit ^48^.

Following auxin treatment of L4 animals, we observed a significant decrease in GRWD-1::degron::GFP and TSR-2::degron::GFP intensity in germ lines, confirming the depletion of both ribosome assembly factors (**Figure 1C-E**). However, RNA-seq analyses revealed no significant reduction in ITS1 and ITS2 pre-rRNA reads, suggesting that the depletion of ribosome assembly factors does not impair rRNA synthesis mediated by RNA Pol I activity (**Figure 1F** and **S1C**, one-tailed Welch’s two-sample *t*-tests; GRWD-1: 1.4-fold average increase in ITS1: *p* = 0.99, 1.4-fold average decrease in ITS2: *p* = 0.16; TSR-2: 1.1-fold average increase in ITS1: *p =* 0.37, no average fold-change in ITS2: *p* = 0.54). Instead, consistent with prior observations involving RPL-3 reduction, the ITS1 intermediate accumulated under GRWD-1 depletion (**Figure S1B**) ^21^. Such accumulation is similar to that reported in other systems, where defects in 60S assembly result in the accumulation of 40S-associated pre-rRNA intermediates containing ITS1 sequences ^49^. RT-qPCR also validated that pre-rRNA synthesis does not decrease upon GRWD-1 depletion (**Figure S1D**, left-tailed paired Student’s *t-*test, ITS1: 0.67, ITS2: *p* = 0.99). Furthermore, the nucleolar morphology in GRWD-1-depleted germ lines appeared vacuolated yet retained a persistent spherical form, similar to the morphology observed when RPL-3 is reduced (**Figure 1H**) ^21^. Polysome profiling further indicates that global translation levels are similarly reduced upon depletion of RPOA-2, GRWD-1, or TSR-2 (**Figure S1E**).

Together, these findings support the conclusion that although RPOA-2 and GRWD-1 depletion both reduce ribosome biogenesis, only impaired rRNA transcription, mediated by a disruption of RNA Pol I activity, disrupts spherical nucleolar structure throughout the germ line. This result highlights a specific requirement for RNA Pol I activity in maintaining nucleolar morphology during oogenesis, independent of downstream ribosome assembly.

### RNA Pol I activity plays a ribosome-biogenesis-independent role in maintaining oocyte production and morphology

To investigate whether germ cells remain viable when RNA Pol I activity and ribosome assembly are reduced, we stained animals with acridine orange to label cell corpses following auxin-inducible depletion of RPOA-2, GRWD-1, or TSR-2. We did not observe any differences in cell corpse abundance in any of our treatments (**Figure S2A-B**), suggesting that apoptosis was not induced during this time frame. Given that germ cells remained viable, we next examined whether germ line maturation was affected upon reductions in Pol I activity or ribosome biogenesis by measuring proximal arm length, the number of proximal oocytes, and the dimensions of the three most proximal oocytes (**Figure 2A-B**). We also normalized proximal arm length to overall body area (**Figure 2C)**, given that the global AID system can reduce body size due to high basal degradation in the absence of auxin, particularly in the GRWD-1 and TSR-2 AID strains ^43^ (**Figure 2A, S2C**).

**Figure 2.**
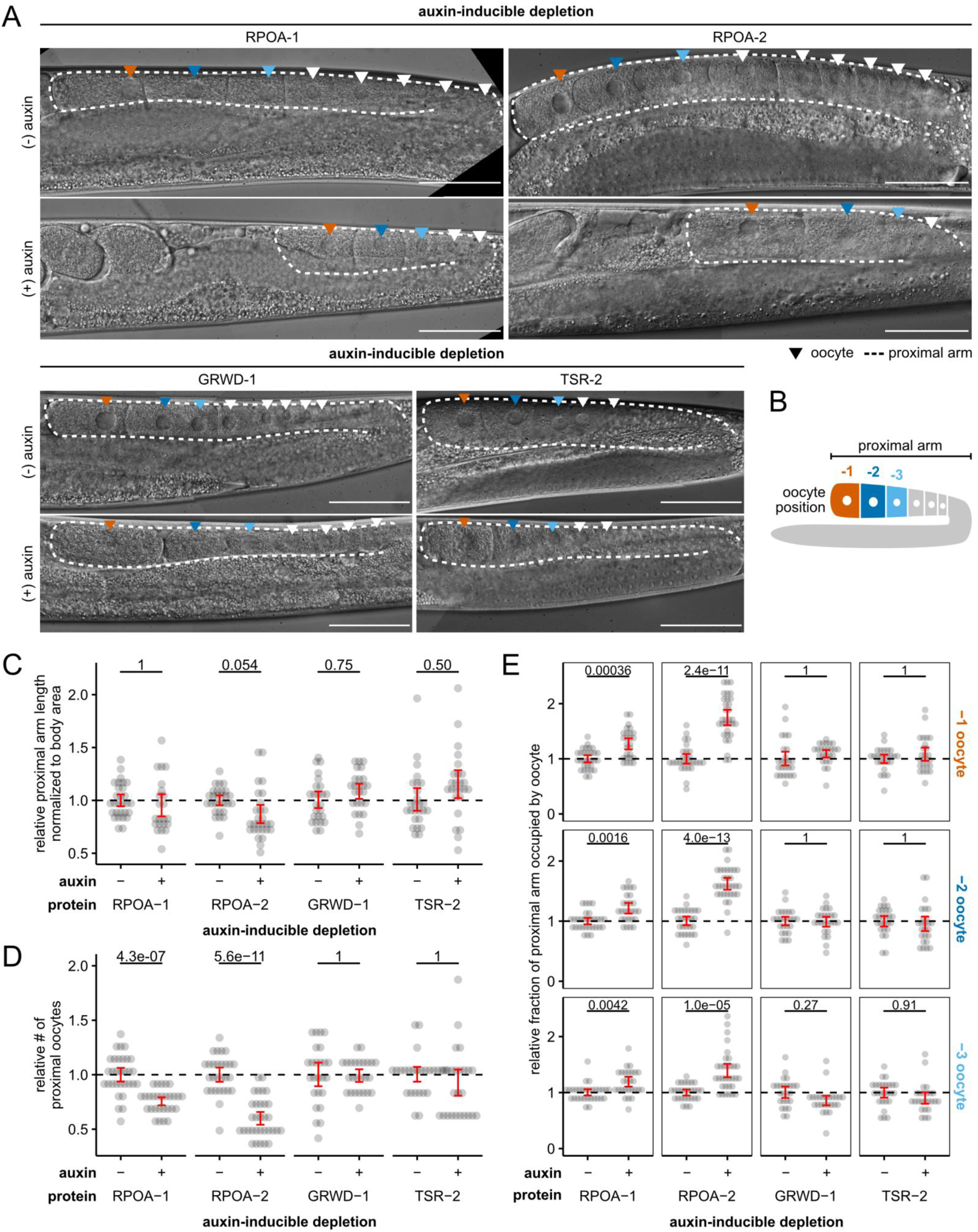
Oocyte morphology and organization are impaired by reduced RNA Pol I activity, independent from RNA Pol I’s role in ribosome production. See also Figure S2. (**A**) Representative DIC images of adult germ lines after auxin-inducible depletion of RPOA-1/RPOA-2/GRWD-1/TSR-2. Dotted lines highlight the proximal arm; triangles point to proximal oocytes. Scale bar = 50 μm. (**B**) Graphical representation of proximal arm length and oocyte positions quantified in (**C-E**). (**C-E**) Proximal arm lengths (**C**), proximal oocyte counts (**D**), and oocyte sizes (**E**) after auxin-inducible depletion of RPOA-1/RPOA-2/GRWD-1/TSR-2. Each point represents a measurement from an individual animal, normalized to the mean of each strain’s control group. Crossbars represent SEM. All comparisons were performed using two-tailed Welch’s two-sample *t*-tests followed by Bonferroni corrections.

Reductions in Pol I activity (mediated by RPOA-1 or RPOA-2 depletion), but not reductions in ribosome assembly (mediated by GRWD-1 or TSR-2 depletion), uniquely reduced the length of the proximal arm (**Figure 2A-B**), diminished the number of proximal oocytes (**Figure 2A** and **2D**), and caused the three most proximal oocytes to occupy a significantly larger fraction of the proximal arm (**Figure 2A** and **2E**). These results suggest that oocyte production and morphology are disrupted specifically by reducing RNA Pol I activity, rather than by impairing ribosome assembly.

### Impaired synapsis reveals a Pol I-dependent sensitivity to nucleolar stress during the mitotic-to-meiotic transition

The observation that reduced RNA Pol I activity, but not impaired ribosome assembly, leads to defects in oocyte maturation prompted us to investigate how each condition affects meiotic progression. Using a fluorescently tagged SYP-3, a component of the synaptonemal complex ^36,37^, we determined whether germ cells adopted characteristic DNA morphology in the mitotic, transition, and meiotic zones. We assessed the appearance of crescent shaped nuclei in the transition zone, where DNA and the nucleolus cluster to opposite nuclear poles, and whether chromosomes in the meiotic zone were repositioned to the nuclear periphery.

Following auxin-inducible depletion of RPOA-2, germ cells entering the mitotic-to-meiotic transition zone showed reduced polarization and SYP-3 assembly, likely reflecting a lack of synaptonemal complex formation; however, the downstream re-localization of synapsed chromosomes to the nuclear periphery in proximal germ cells remained unaffected (**Figure 3A, S3)**.

**Figure 3.**
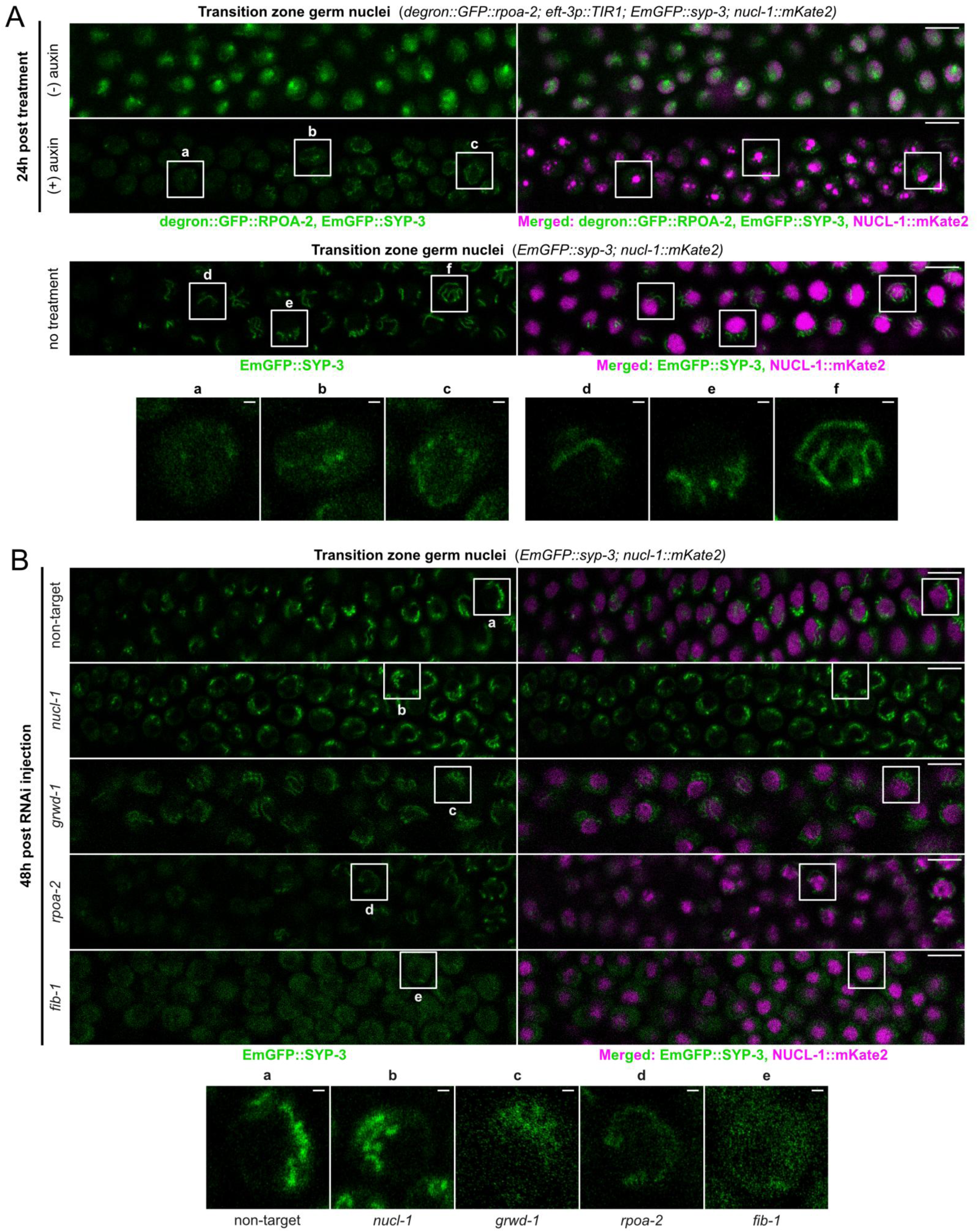
Synaptonemal complex formation during the mitotic-to-meiotic germ cell transition is dependent on Pol I activity, but not other ribosome processing steps. See also Figure S3 and S4. **(A)** Confocal images of transition zone nuclei expressing degron::GFP::RPOA-2, EmGFP::SYP-3, and NUCL-1::mKate2 following control or auxin-inducible depletion of RPOA-2 (top two rows). Additionally, we show non-treated nuclei from a wild-type genetic background (middle row) and zoomed-in nuclei from RPOA-2-depleted and wild-type conditions (bottom row, labeled a-e). Scale bar = 5 μm (transition zone) and 0.5 μm (zoomed-in nuclei). **(B)** Confocal images of transition zone nuclei expressing EmGFP::SYP-3 and NUCL-1::mKate2 48 hours post RNAi knockdown of a non-target locus (*wrmScarlet*), *nucl-1*, *grwd-1*, *rpoa-2*, and *fib-1* (top), as well as zoomed-in nuclei from each condition (bottom). Scale bar = 5 μm (transition zone) and 0.5 μm (zoomed-in nuclei). All images were taken with identical laser intensity; due to decreased SYP-3 expression under knockdown of *grwd-1, rpoa-2,* and *fib-1*, we have increased the brightness of these conditions to be more comparable to wild type.

These differences raised the possibility that nucleolar integrity is critical for proper chromosome morphology during meiotic entry. Specifically, in transition zone nuclei, the nucleolus and chromosomes segregate to opposite poles, likely to facilitate homolog pairing by bringing homologous chromosomes into proximity ^50–52^. To determine whether perturbing nucleolar structure or activity, rather than bulk ribosome synthesis, affects this characteristic crescent shaped chromosome polarization, we depleted key ribosome synthesis and nucleolar factors by RNA interference (RNAi). We first validated RNAi efficacy by injecting double-stranded RNA targeting *rpoa-2, grwd-1, fib-1* (human *FBL* ortholog), and *nucl-1*, and monitored protein depletion using fluorescent tagged reporters. All four proteins were effectively depleted to near-background levels within 48 hours (**Figure S4A-D**). We then used these validated RNAi conditions on an EmGFP::SYP-3 and NUCL-1::mKate2 reporter strain to examine synaptonemal complex assembly and chromatin architecture throughout meiotic prophase (**Figure 3B**, **S4E**). Depletion of *rpoa-2* and *fib-1* resulted in similar germline deficiencies, including less transition zone polarization **(Figure 3B, S4E)**, a reduction of transition zone SYP-3 aggregation (indicative of impaired synapsis) **(Figure 3B, S4E),** and a reduction of proximal oocytes **(Figure S4F, 2A).** In contrast, *nucl-1* knockdown resulted in no detectable defects in either chromosome polarization or synapsis. Similarly, although knockdown of *grwd-1* decreased overall SYP-3 intensity, SYP-3 aggregation and transition zone polarization remained largely comparable to non-target RNAi injected gonads (**Figure 3B**, **S4E**). In addition, we did not observe any differences in the redistribution of SYP-3 to the nuclear periphery in late pachytene/post transition zone nuclei in any of the knockdowns (**Figure S4E**), likely suggesting that newly transitioning nuclei are receptive to nucleolar stress.

Together, these results support that nuclei entering the mitotic-to-meiotic transition zone are sensitive to nucleolar functions dependent on RNA Pol I activity (RPOA-2 and FIB-1) rather than downstream ribosome processing steps (NUCL-1 and GRWD-1), supported by Pol-I dependent defects in transition zone polarization and synaptonemal complex assembly.

### Autosomal oogenesis gene promoters are more accessible after RNA Pol I depletion

Given that RNA Pol I levels were associated with germline chromosome abnormalities, we next investigated whether specific genomic regions were preferentially affected by a reduction of RNA Pol I activity compared to reduced ribosome assembly. We performed ATAC-seq on dissected gonads from control and RPOA-2- and GRWD-1-depleted young adults. In RPOA-2-depleted germ lines, 1,616 genomic regions were significantly more accessible (SMA), whereas only 99 regions were significantly less accessible (DESeq2, Benjamini-Hochberg adjusted *p* < 0.05). In GRWD-1-depleted germ lines, 53 regions were more accessible and only one region showed reduced accessibility (DESeq2, Benjamini-Hochberg adjusted *p* < 0.05). Additionally, 41 regions were SMA in both RPOA-2- and GRWD-1-depleted gonads (**Figure 4A**).

**Figure 4.**
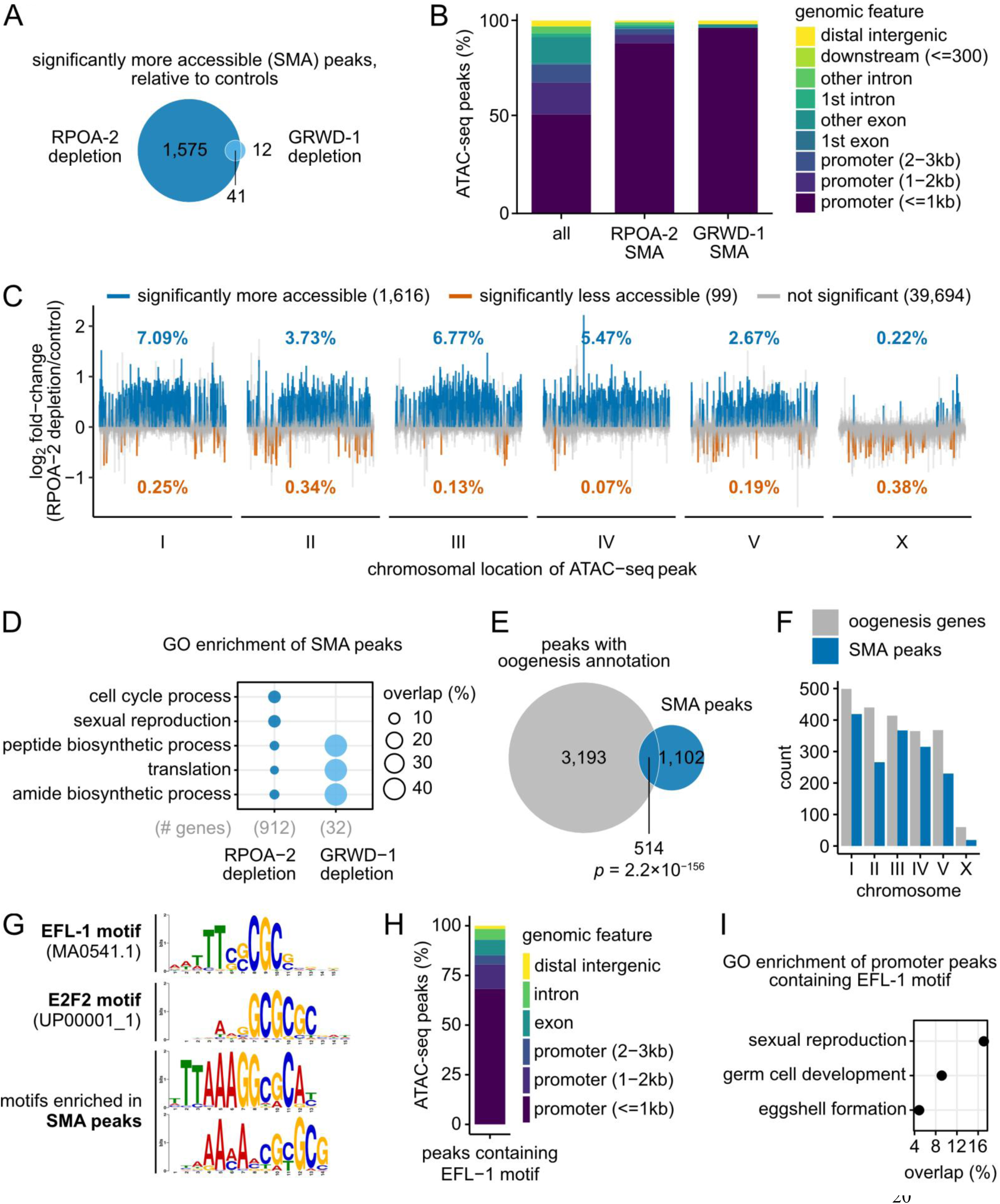
Disrupting RNA Pol I activity prematurely primes autosomal promoters in an accessible state associated with oogenesis. See also Figure S5. (**A**) Venn diagram comparing the number of significantly more accessible (SMA) regions following RPOA-2 depletion versus GRWD-1 depletion, based on an adjusted *p* < 0.05. (**B**) Genomic feature annotations of ATAC-seq peaks. SMA regions after RPOA-2 or GRWD-1 depletion are predominantly located within 1 kb of the nearest promoter. (**C**) Log_2_ fold-change estimates of chromatin accessibility based on gonadal ATAC-seq data from at least three biological replicates per condition, each composed of 20 gonads. Each segment along the x-axis represents a genomic region. The number of differentially accessible regions are noted in the plot legend based on an adjusted *p* < 0.05. Percentages represent the proportion of differentially accessible peaks relative to the total number of called peaks per chromosome. (**D**) Enrichment of gene ontology (GO) biological processes in genes exhibiting increased accessibility after depletion of RPOA-2 or GRWD-1 (Hypergeometric test, FDR < 0.001). The term “gene” refers to those associated with the nearest genomic feature annotation of each peak. (**E**) Overlap between peaks that are SMA after RPOA-2 depletion and peaks near genes involved in oogenesis ^53^ (Hypergeometric test). (**F**) Chromosomal distribution of genes involved in oogenesis ^53^ and SMA peaks after RPOA-2 depletion. (**G**) Alignment of the *C. elegans* EFL-1-binding site, human E2F-binding site, and the two motifs enriched in SMA peaks after RPOA-2 depletion. (**H**) Genomic feature annotations of peaks containing an EFL-1-binding site. (**I**) GO enrichment analysis for biological processes among genes with promoter peaks that contain an EFL-1-binding site (Hypergeometric test, FDR < 0.05).

To functionally interpret these changes, we annotated each ATAC-seq peak to its nearest genomic feature and found that more than 80% of SMA peaks in either RPOA-2- or GRWD-1-depleted germ lines were located within 1 kb of the promoter of the nearest gene, compared to only 50% in background peaks (**Figure 4B**). Moreover, SMA regions in RPOA-2-depleted germ lines were concentrated on autosomes (**Figure 4C**), with ∼2-7% of peaks per autosome showing increased accessibility, compared to only ∼0.2% on the X chromosome (**Figure 4C**). In GRWD-1-depleted germ lines, SMA regions on the X chromosome were also less prominent, though to a lesser extent (**Figure S5A**). Overall, these results indicate that nucleolar disruption caused by reduced RNA Pol I activity leads to a more pronounced increase in chromatin accessibility at autosomal promoters, emphasizing the role of nucleolar integrity in genome organization.

### RNA Pol I-driven nucleolar disruption prematurely primes oogenesis promoters in an accessible state

Considering that RNA Pol I levels were linked to increased chromatin accessibility along the promoters of autosomal genes, we asked whether the genes affected by these accessibility changes contribute to the defective oocyte maturation that emerges specifically under reduced RNA Pol I activity. To examine this, we performed a differential gene ontology (GO) analysis of biological processes enriched in genes with increased accessibility following RPOA-2 or GRWD-1 depletion. Regions with increased accessibility following a reduction of RNA Pol I activity were enriched for sexual reproduction and cell cycle processes, including meiotic cell cycle, chromosome organization, and oogenesis (**Figure 4D** and **Table S2**, Hypergeometric test, FDR < 0.001). Additionally, RPOA-2 depletion uniquely increased the accessibility of genes that function both in germ/oocyte development and translation (e.g., *daz-1, fbf-2, fog-1, gld-1*, *glp-4*, *larp-1, mex-3, oma-1*, *oma-2*) (**Figure 4D** and **Table S2**, Hypergeometric test, FDR < 0.001). In contrast, depleting GRWD-1, which impairs ribosome assembly without disrupting spherical nucleolar structure, did not show enrichment of sexual reproduction processes in more accessible chromatin regions. Instead, genes related to translation, mostly consisting of ribosomal protein genes, mitochondrial-ribosome-related genes, and translation initiation/termination factors (e.g., *rps-0, rps-9, rps-17, rps-27, rpl-9, rpl-18, rpl-25.2, rpl-28, inf-1, larp-5, rack-1, W01D2.1*) showed increased accessibility under both RPOA-2 and GRWD-1 depletion, consistent with adaptive response to reduced ribosome production. These results suggest that although decreased ribosome production increases accessibility at translation related loci in both conditions, reducing RNA Pol I activity uniquely increases the accessibility of oogenesis genes, potentially contributing to the defective oogenesis phenotype (**Figure 2**).

We further investigated the extent to which oogenesis genes were affected by RNA Pol I depletion using a set of 2,177 genes with germline-enriched expression in feminized *fem-1(lf)* mutants ^53^. Of the 3,707 peaks annotated to these oogenesis genes, we detected a significant enrichment within regions that became significantly more accessible after RPOA-2 depletion (**Figure 4E**, Hypergeometric test, *p* = 2.2×10^−156^). Consistent with this observation, the autosomal bias of oogenesis genes parallels the autosomal bias in accessibility observed for RPOA-2-depleted gonads (**Figure 4F**).

To assess potential regulators of these accessibility changes, we searched for enriched transcription factor (TF) binding motifs that gained accessibility after RPOA-2 depletion. Compared to ungapped sequence motifs present in background peaks, we identified eight motifs significantly enriched in the accessible peaks of RPOA-2-depleted germ cells (**Table S3**, STREME, *p* < 0.05, *E* < 0.05). We aligned the eight motifs enriched in RPOA-2-depleted germ lines against the 842 TF motifs present in the TFBSshape database and identified 143 unique TF binding motif matches in peaks with increased accessibility after RPOA-2 depletion (Tomtom, *p* < 0.05, *E* < 8.42) ^54^. After manual curation to combine redundant query motifs and their repeated matches, we found relatively few matches with visibly cohesive alignments to their query motifs. Notably, we identified four total hits to human E2F-binding sites from the enriched RTGCGCCTTTAAA and CGCRCGKTKTTTRN motifs (**Figure 4G**, **Table S3**), which are orthologs to the oogenesis-promoting *C. elegans* transcription factor EFL-1 ^55^. Additionally, the CGCRCGKTKTTTRN motif aligned to the yeast Rsc3- and Rsc30-binding sites, which regulate ribosomal protein gene expression ^56^.

Although there was no enrichment of motifs for accessible regions following GRWD-1 depletion, we corroborated that the EFL-1-like binding motif was specifically more accessible following RPOA-2 depletion, but not after GRWD-1 depletion. We identified 242 ATAC-seq peaks containing a significant match to the EFL-1 motif, predominantly in promoters less than 1 kb from the nearest gene (**Figure 4H**, FIMO, *p* < 0.0001). Genes with promoter peaks containing an EFL-1 motif are involved in processes such as sexual reproduction, germ cell development, and eggshell formation (**Figure 4I**, Hypergeometric test, FDR < 0.05). Lastly, genes with a promoter peak containing the EFL-1 motif were not remodeled after reducing ribosome assembly but did show increased accessibility after reducing RNA Pol I activity (**Figure S5F**).

We next examined whether these accessibility changes affected steady-state transcript levels by analyzing RNA-seq data from dissected gonads after RPOA-2 or GRWD-1-depletion. We identified 188 genes with increased mRNA levels and 177 genes with reduced mRNA levels upon RPOA-2 depletion (**Figure S5B**, Benjamini-Hochberg adjusted *p* < 0.05). Similarly, GRWD-1 depletion led to increased mRNA levels in 287 genes and decreased levels in 235 genes (**Figure S5B**, Benjamini-Hochberg adjusted *p* < 0.05). We did not observe significant functional GO enrichment in the over-expressed genes unique to RPOA-2 depletion (**Table S5**, Hypergeometric test, FDR < 0.001). Both conditions shared 57 over-expressed genes and 87 under-expressed genes with a shared enrichment of under-expressed genes related to translation (**Figure S3C-D**, Hypergeometric test, FDR < 0.001), mirroring patterns observed in chromatin accessibility. Moreover, we found no overall correlation between chromatin accessibility and mRNA levels for all genes or for those that were differentially accessible and differentially expressed (**Figure S5E**, *p* > 0.05).

Given the enrichment of a putative repressive Rsc30-like motif in accessible regions following RPOA-2 depletion, we hypothesized that translation-related genes would be under-expressed. Indeed, many ribosomal protein genes are significantly reduced after RPOA-2 depletion, consistent with negative regulation (**Figure S5G**, one-tailed one-sample *t-*test, alternative hypothesis: average log_2_ fold-change < 0, *p* = 0.00020). Such coordinated repression can be explained by previously described feedback mechanisms that couple ribosomal protein gene expression directly to rRNA transcription. Specifically, the transcriptional activator Ifh1 interacts with the rRNA processing factor Utp22 to synchronize ribosomal protein gene transcription with rRNA synthesis ^57^. Under conditions of impaired rRNA synthesis, as occurs with RPOA-2 depletion, Utp22 is no longer sequestered by rRNA and thus becomes available to bind ribosomal protein gene promoters, leading to transcriptional repression. In contrast, depletion of GRWD-1 disrupts assembly of the 60S ribosomal subunit without directly impairing rRNA transcription (**Figure 1F, S1C, S1D**). As a result, the Utp22-mediated feedback inhibition of ribosomal protein genes is likely not triggered, and instead these genes become overexpressed as a compensatory response to defective ribosome assembly (**Figure S5D**).

In contrast, genes related to sexual reproduction did not exhibit increased transcript levels, despite increased chromatin accessibility and the enrichment of a potential oogenesis-promoting transcription factor motif (**Figure S5G**, one-tailed one-sample *t*-test, alternative hypothesis: average log_2_ fold-change > 0, *p* = 0.16). In agreement, genes with a promoter peak containing an EFL-1 motif did not significantly increase mRNA expression after RPOA-2 depletion (**Figure S5F**, one-tailed one-sample *t*-test, alternative hypothesis: log_2_ fold-change > 0, *p* = 0.14). Thus, although oogenesis promoters become more accessible, mRNA levels remain unchanged, reflecting an oogenesis-specific chromatin state that is primed for but does not undergo productive transcription.

Together, these results suggest that decreasing RNA Pol I activity triggers nucleosome remodeling to reduce the expression of ribosomal proteins, potentially conserving resources in response to limited rRNA availability for ribosome biogenesis. However, this reduction in RNA Pol I activity also leads to nucleosome remodeling that increases the accessibility of oogenesis-related genes and oogenesis-promoting EFL-1 binding sites, potentially contributing to a defective oocyte maturation phenotype (**Figure 2**).

### Increases in chromatin accessibility upon Pol I depletion are accompanied by oogenesis-specific histone remodeling

Throughout oogenesis, germ cells undergo extensive chromatin remodeling to prepare for meiotic chromosome condensation and subsequent zygotic genome activation (ZGA). One hallmark of this transition is the redistribution of H3K4me3 from gene bodies to promoters in mature oocytes ^4^. Given that RPOA-2 depletion increases the accessibility of oogenesis-related regions, we asked whether these loci show typical signatures of H3K4me3 remodeling in maturing wild-type oocytes.

Genes undergoing H3K4me3 remodeling during oogenesis can be grouped into three clusters: Cluster 1 (strong H3K4me3 marks in both promoters and gene bodies), Cluster 2 (strong promoter-specific H3K4me3), and Cluster 3 (weaker promoter-specific H3K4me3) ^4^. When we compared chromatin accessibility near genes in these clusters, we found that accessibility near oogenesis-specific H3K4me3 remodeled genes significantly increased after RPOA-2 depletion relative to a random set of regions (**Figure 5A**, Bonferroni-corrected one-tailed Welch’s two-sample *t*-tests, *p* < 0.05). Consistent with the germ-to-oocyte H3K4me3 remodeling pattern, Cluster 1 genes showed increased chromatin accessibility along both promoters and gene bodies (**Figure 5B**, top). Cluster 2 genes exhibited increased accessibility mainly in promoters, while Cluster 3 genes displayed weaker promoter specific increases following RPOA-2 depletion (**Figure 5B**, middle and bottom).

**Figure 5.**
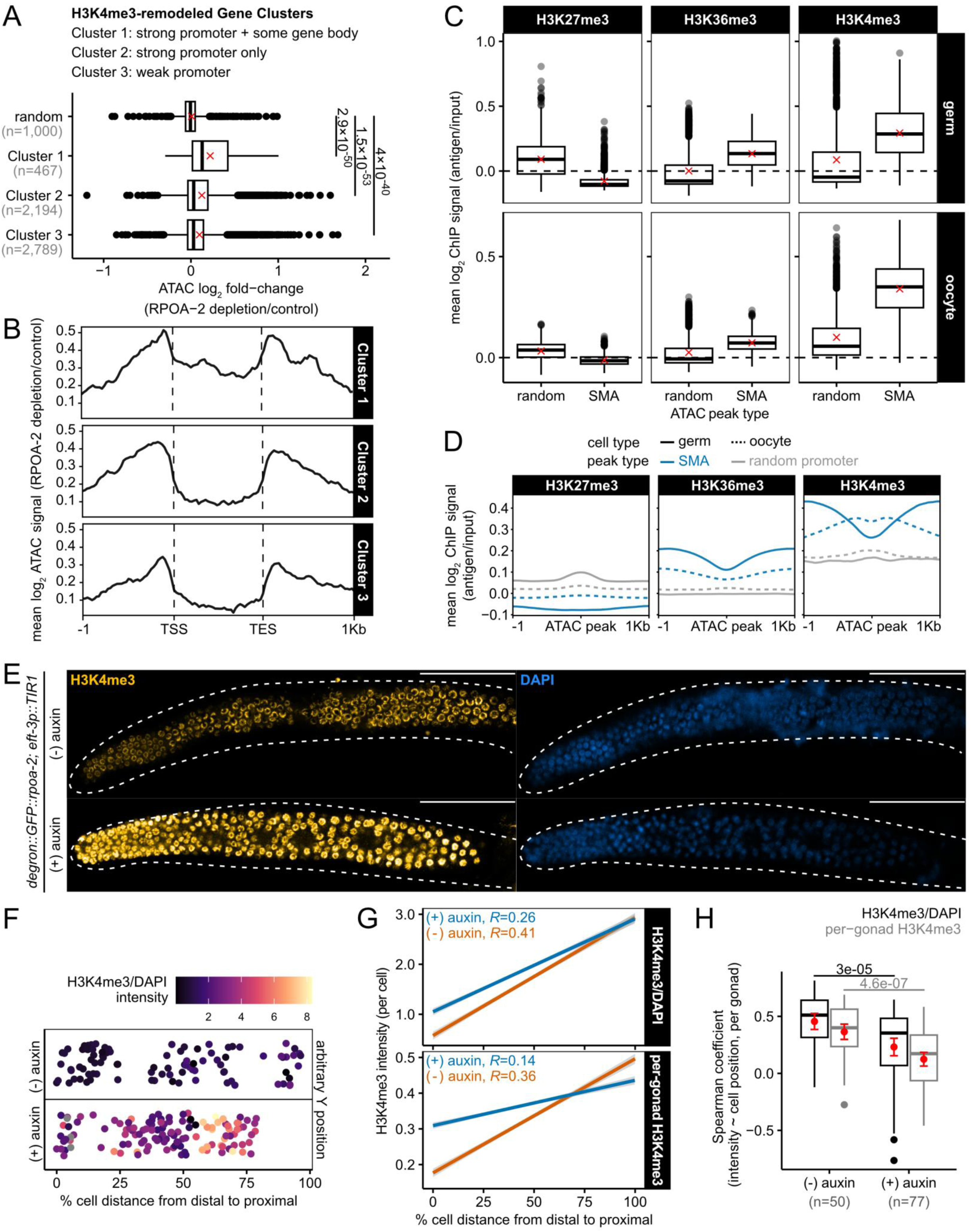
Chromatin regions that become more accessible after reducing RNA Pol I activity display patterns of H3K4me3 remodeling characteristic of the germ-to-oocyte transition. See also Figure S6. (**A**) Log_2_ fold-change estimates of chromatin accessibility after RPOA-2 depletion in peaks annotated to genes that undergo H3K4me3 remodeling during oogenesis, compared to a random subset of peaks. The number of peaks in each set is specified by *n*, and means are denoted by an *x*. Statistical comparisons were performed using right-tailed Welch’s two-sample *t*-tests with Bonferroni corrections. (**B**) Metagene analysis of mean log_2_ ATAC signal (RPOA-2 depletion/control) within 1 kb of genes that undergo H3K4me3 remodeling during oogenesis. In gonads with reduced RNA Pol I activity, the shape of chromatin accessibility signal within each cluster mirrors the shape of H3K4me3 remodeling associated with each gene cluster. (**C**) Distribution of average H3K27me3, H3K36me3, and H3K4me3 ChIP signals from wild-type germ cells or oocytes within ATAC-seq peaks that are significantly more accessible (SMA) after RPOA-2 depletion, compared to signals within an identically-sized random subset of regions. Means are denoted by an *x*. (**D**) Metagene plot displaying the average H3K27me3, H3K36me3, and H3K4me3 ChIP coverage from wild-type germ cells or oocytes within 1 kb of genomic regions that are SMA after RPOA-2 depletion, compared to coverage within an identically-sized random subset of promoter ATAC-seq peaks that are not SMA. (**E**) Representative 3-D confocal projections of adult gonad arms with H3K4me3 immunostaining and DAPI after control treatment or RPOA-2 depletion. Dotted lines represent gonad boundaries determined from T-PMT. Scale bar = 50 μm. (**F**) Using the example gonads from (E), we show germ cell position along a linear gonad axis following implementation of a gonad linearization algorithm (see Methods). Each point represents a germ cell detection; however, note that not all gonadal cells are detected by the algorithm. The distal tip is represented by zero on the gonad axis, whereas the most proximal tip of the gonad is variable based on biological size, gonad disruption during immunostaining, or gonad placement on Z-plane. H3K4me3 intensity measurements are normalized to DAPI to account for differences in permeability across gonads. (**G**) Linear relationship between germ cell position along the gonadal axis (from distal to proximal end) and H3K4me3 signal intensity (relative to DAPI intensity or scaled per-gonad), aggregated across all gonads. Spearman correlation coefficients are represented on the plot. Linear models and overall correlations are based on data from 3,933 control cell detections and 8,134 depletion cell detections from 50 and 77 gonads, respectively. (**H**) Distribution of Spearman coefficients estimating per-gonad positive correlations between cell position on the gonad axis and H3K4me3 intensity (relative to DAPI intensity or scaled per-gonad). Mean and standard error are represented by red points and crossbars. The number of gonads analyzed per condition are represented by *n*. A one-tailed Welch’s two-sample *t*-test suggests that on average, control gonads have a higher magnitude correlation than gonads with reduced RNA Pol I activity (Benjamini-Hochberg adjusted *p*-values).

Although reducing RNA Pol I activity shows signatures of H3K4me3 remodeling, changes in chromatin accessibility can also reflect the remodeling of other histone marks. We compared the abundance of three such histone marks—remodeled active H3K4me3, non-remodeled active H3K36me3, and non-remodeled repressive H3K27me3—within regions that became more accessible after RPOA-2 depletion, using available ChIP-seq data from wild-type germ cells and oocytes ^58–60^. In wild-type germ cells, H3K27me3 marks were less abundant in SMA regions compared to random regions (**Figure 5C**, top left), while active marks (H3K36me3 and H3K4me3) were enriched in SMA regions (**Figure 5C**, top middle and right). In wild-type oocytes, we expected highly accessible regions after RPOA-2 depletion to show specific enrichment for H3K4me3, reflecting germ-to-oocyte histone remodeling. In agreement, H3K4me3 marks were enriched in SMA regions relative to random regions (**Figure 5C**, bottom right), while H3K27me3 and H3K36me3 signals were weak in both accessible and random regions (**Figure 5C**, bottom left and middle).

Since more than 80% of SMA peaks are near promoters (**Figure 4B**), and H3K4me3 marks shift to promoters in oocytes, we hypothesized that H3K4me3 signal would peak at the center of SMA regions in wild-type oocytes. In contrast, we expected H3K4me3 in wild-type germ cells to decrease around SMA regions, since H3K4me3 is typically present along gene bodies in germ cells. This pattern was confirmed, with H3K4me3 signal peaking in SMA regions relative to a random set of promoters in wild-type oocytes (**Figure 5D**, right) and decreasing around SMA regions in wild-type germ cells. H3K27me3 and H3K36me3 signals did not notably peak near SMA regions in either germ cells or oocytes (**Figure 5D**, left and middle). Our findings support that reducing RNA Pol I activity throughout the germ line specifically increases the accessibility of promoters that are remodeled during the germ-to-oocyte transition through H3K4me3 deposition, suggesting that the spatiotemporal regulation of H3K4 methylation is misregulated under RNA Pol I-mediated nucleolar disruption.

In wild-type animals, H3K4 methylation progressively increases towards the proximal region of the gonad, and this spatial regulation of H3K4 methylation is essential for proper germ line maturation ^12,61^. Given that RPOA-2 depletion increases the accessibility of oogenesis-associated H3K4me3 regions and specifically disrupts oogenesis, we anticipated that the spatial regulation of H3K4 methylation would be altered in response to RPOA-2 depletion, likely through premature H3K4me3 deposition in accordance with increased chromatin accessibility. To investigate this, we performed gonadal H3K4me3 immunostaining and cell-segmentation image analysis to assess how reduced RNA Pol I activity affects changes in the deposition of H3K4me3 from the distal to proximal end of the gonad (**Figure 5E**, *n =* 50 control gonads with 3,933 cells, *n* = 77 depletion gonads with 8,134 cells). Individual germ cell positions were projected onto a linear gonad axis to evaluate how H3K4me3 immunostaining intensity correlated with cell position along the distal-proximal gonad axis (termed cell gonad position from here on) (**Figure 5F**, see Methods).

As expected, an aggregate analysis of cells from all control gonads revealed a significant gradual increase in H3K4 methylation along the distal-proximal axis, evidenced both by H3K4me3/DAPI normalized intensity (**Figure 5G, top**, Spearman coefficient = 0.41) and by per-gonad scaled H3K4me3 intensity (**Figure 5G, bottom**, Spearman coefficient = 0.36). Similarly, we also analyzed this positive H3K4me3 gradient within individual gonads and observed that the majority (84%) of control gonads had significant correlations between cell gonad position and normalized H3K4me3/DAPI intensity (**Figure 5H**, mean Spearman coefficient: 0.46±0.03, Benjamini-Hochberg adjusted *p*-value < 0.05), and also per-gonad scaled H3K4me3 intensity (**Figure 5H**, mean Spearman coefficient: 0.36±0.03, 78% with Benjamini-Hochberg adjusted *p*-value < 0.05). These results support that typical oogenesis is consistently associated with a moderate but important positive H3K4me3 gradient.

Under reduced RNA Pol I activity, the aggregated correlation between cell gonad position and H3K4me3 was reduced by approximately half, both relative to normalized H3K4me3/DAPI and per-gonad scaled H3K4me3 intensity (**Figure 5G**, Spearman coefficient = 0.26 and 0.14 for H3K4me3/DAPI and per-gonad scaled H3K4me3, respectively). Additionally, only 64% of RPOA-2-depleted gonads had significant, but weaker, correlations between cell gonad position and normalized H3K4me3/DAPI intensity (**Figure 5H**, mean Spearman coefficient: 0.23±0.04, Benjamini-Hochberg adjusted *p*-value < 0.05) and also per-gonad scaled H3K4me3 intensity (**Figure 5H**, mean Spearman coefficient: 0.12±0.03, 52% with Benjamini-Hochberg adjusted *p*-value < 0.05).

Lastly, compared to control gonads, RPOA-2-depleted gonads had a significant reduction in average correlation between cell gonad position and H3K4me3 (**Figure 5H**, one-tailed Welch’s two-sample *t*-test, H3K4me3/DAPI: Benjamini-Hochberg adjusted *p*-value = 3.0×10^-5^, per-gonad scaled H3K4me3: Benjamini-Hochberg adjusted *p*-value = 4.6×10^-7^). Interestingly, this misregulation of H3K4me3 deposition could be explained by increased H3K4me3 in the distal region (**Figure 5G, top**).

Given RPOA-2 depletion specifically disrupts the spatiotemporal regulation of H3K4 methylation, premature and aberrant deposition of H3K4me3 likely interferes with synapsis and meiotic chromosome organization. Indeed, previous studies have established that disrupted H3K4 methylation dynamics strongly correlate with recombination abnormalities, and defects in synaptonemal assembly and stability ^62,63^. Consistent with these findings, we also observed synapsis defects indicated by altered SYP-3 localization, following RPOA-2 depletion (**Figure 3A-B, S4**).

To further dissect the mechanistic basis underlying these defects, we asked whether nucleolar disruption and associated chromatin alterations activate the meiotic DNA damage checkpoint kinases ATM-1 and ATL-1. Immunostaining for phosphorylated serine/threonine residues, a hallmark of ATM/ATR checkpoint activation, revealed visibly increased checkpoint signaling specifically upon RPOA-2 depletion, but not after GRWD-1 depletion, suggesting checkpoint activation is specific to RNA Pol I mediated nucleolar stress (**Figure S6A**).

ATM/ATR activation is known to trigger CEP-1 (p53) mediated apoptosis in the presence of extensive DNA damage or unresolved meiotic recombination intermediates. Thus, we asked whether increased ATM/ATR phosphorylation following RPOA-2 depletion led to activation of CEP-1/p53. RNAi experiments targeting *rpoa-2* or *grwd-1* in a CEP-1::GFP reporter strain showed no visible increase in CEP-1 expression compared to control RNAi (**Figure S6B**). These data indicate that the meiotic chromosome defects and associated checkpoint activation induced by RNA Pol I depletion occur independently of apoptosis, consistent with acridine orange staining results, which revealed no significant increase in apoptotic cells (**Figure S2A, B**). Instead, these results suggest that ATM/ATR checkpoint activation occurs in conjunction with synapsis defects.

Given the proposed role of SET-2 (the *C. elegans* homolog of SET1 responsible for catalyzing H3K4 methylation) in preserving genome integrity and regulating chromatin compaction in the germ line ^63,64^, we hypothesized that SET-2 might mediate the aberrant H3K4 methylation observed following RPOA-2 depletion. We further reasoned that loss of SET-2 function might rescue or alter the oogenesis defects and abnormal methylation patterns seen after RPOA-2 depletion. To test this hypothesis, we crossed the *set-2(ok592)* mutant into the RPOA-2 AID strain. However, RPOA-2 depletion in the *set-2(ok592)* mutant background still resulted in a reduced number of enlarged oocytes (**Figure S6C**), indicating that SET-2 does not significantly influence the RNA Pol I dependent oogenesis defects. Furthermore, H3K4me3 immunostaining revealed persistently increased H3K4 trimethylation upon RPOA-2 depletion in the absence of SET-2 function (*set-2(ok592*)), suggesting that SET-2 is not the methyltransferase responsible for the aberrant methylation triggered by loss of RPOA-2 (**Figure S6D**).

Together, our data support a model (**Figure 6**) in which RNA Pol I–dependent nucleolar integrity is essential to maintain the proper spatiotemporal regulation of H3K4 methylation during oogenesis. Specifically, reducing RNA Pol I activity results in premature and ectopic deposition of H3K4me3 at the promoters of oogenesis-related genes, leading to abnormal chromatin accessibility patterns. These aberrant chromatin states likely interfere with synaptonemal complex formation and homologous recombination, triggering sustained ATM/ATR checkpoint activation independent of CEP-1/p53 mediated apoptosis. Rather than apoptosis, this checkpoint activation likely leads to meiotic cell-cycle perturbation, suggesting an epigenetic feedback loop that safeguards germline genome integrity upon nucleolar disruption and associated chromatin misregulation (**Figure 6**).

**Figure 6.**
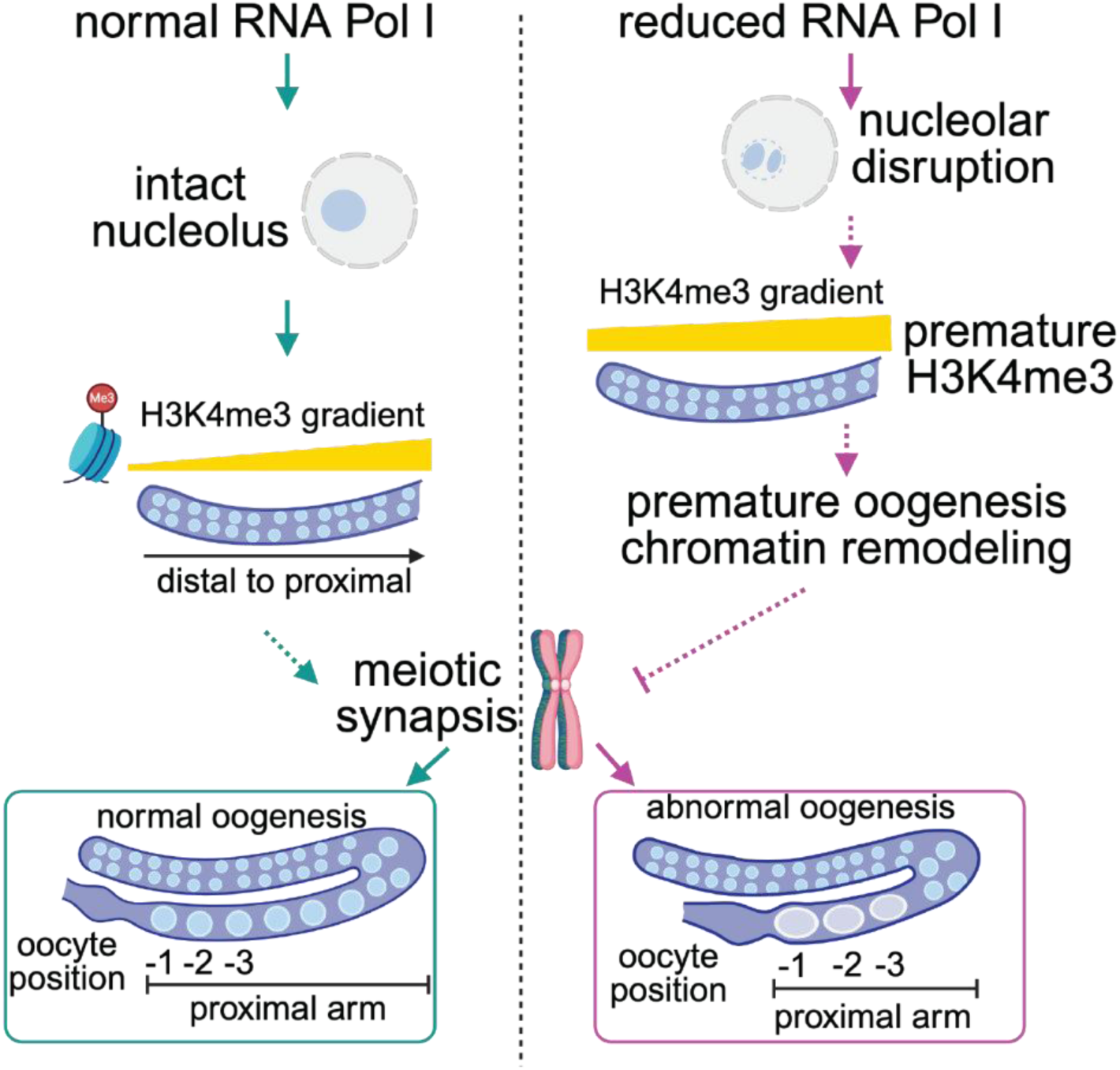
Schematic model illustrating the non-ribosomal function of RNA Pol I during *C. elegans* oogenesis. Under normal conditions (left), RNA Pol I activity maintains nucleolar integrity, which supports the establishment of a proper H3K4me3 gradient from the distal to proximal germ line. This gradient facilitates meiotic synapsis and the progression of oogenesis, resulting in normal oocyte formation. When RNA Pol I activity is reduced (right), nucleolar disruption leads to premature and ectopic H3K4me3 deposition and aberrant oogenesis chromatin remodeling. This dysregulation interferes with the mitotic-to-meiotic transition, resulting in decreased synapsis that leads to abnormal oogenesis, characterized by fewer and enlarged oocytes. The model proposes an epigenetic feedback mechanism that links nucleolar function to germline genome integrity. Created in BioRender. https://BioRender.com/oq1fz8k.

## DISCUSSION

In this study, we investigated non-ribosomal functions of the nucleolus during oogenesis using separation-of-function strains in *Caenorhabditis elegans*. By leveraging the auxin-inducible degron (AID) system, we selectively depleted catalytic subunits of RNA Polymerase I (RPOA-1 and RPOA-2) and two ribosome assembly factors (GRWD-1 and TSR-2) in the germ line. This approach enabled us to distinguish between the effects of nucleolar disruption and reduced ribosome biogenesis, revealing that germ cell nucleoli actively regulate meiotic progression through epigenetic mechanisms independent of ribosome production. Our findings uncover a previously unrecognized role for RNA Pol I activity and nucleolar integrity in oogenesis beyond ribosome production.

### Differential impacts of nucleolar versus ribosomal stress on oogenesis and meiotic synapsis

A key insight from our work is the distinct impact of nucleolar versus ribosomal stress on germ cell differentiation. Near-complete reduction of ribosome assembly had minimal consequences for oocyte production (**Figure 2**), suggesting that the oogenesis program, including meiotic synapsis and histone remodeling, is not heavily dependent on ribosome biogenesis (**Figure 3-4**). In contrast, disruption of nucleolar structure through near-complete reduction of RNA Pol I activity leads to significant oogenesis and synapsis defects (**Figure 2-3**). These results suggest that nucleolar integrity, rather than downstream capacity for protein synthesis, is essential for proper germ cell progression through oogenesis.

Our previous work highlights the importance of ribosome loading in maturing oocytes to support embryogenesis in *C. elegans* ^22^. However, our current results show that ribosome biogenesis is not critical for oogenesis itself, at least within the timeframe of our investigation. Furthermore, we previously demonstrated that somatic chromatin reorganization and developmental progression beyond the L2 stage are similarly affected by either nucleolar or ribosomal stress, indicating a uniform response in somatic tissues ^43,65^. Here, however, germ cells entering meiosis exhibit a unique sensitivity to nucleolar disruption, suggesting that meiotic cells, with their distinct chromatin remodeling requirements, are particularly vulnerable to changes in nucleolar homeostasis.

One explanation may be that meiotic germ cells, which are relatively quiescent in their transcription and translation activities, do not strongly depend on ongoing ribosome production. However, their reliance on large–scale chromosome restructuring places them at risk when nucleolar integrity is compromised; several lines of evidence in other organisms and tissues suggest that nucleolar integrity is linked to genome organization and stress sensing ^66–71^. In female gametogenesis in particular, intact nucleolar structure has been shown to have several implications for successful oogenesis unrelated to ribosome production. These functions include safeguarding genomic stability by shielding repetitive rDNA loci from recombination, preserving satellite repeat sequences, and sequestering transcriptional silencing factors, all of which are crucial for successful oogenesis ^23–25^. Thus, given the potential consequences of aberrant nucleolar structure in maturing germ cells, it is likely that nucleolar stress-sensing mechanisms play a role in regulating meiotic progression.

In support of this view, reductions in ribosome assembly, which maintains a spherical nucleolar structure, does not noticeably impair the mitotic-to-meiotic transition as assessed by SYP-3 localization in the transition zone and pachytene stages (**Figure 3**). In contrast, depletion of RNA Pol I activity, accompanied by nucleolar disruption, impairs synapsis and significantly reduced oocyte production (**Figure 2-3**). Intriguingly, depletion of fibrillarin, a structural nucleolar protein, similarly inhibited meiotic synapsis and reduced oogenesis, suggesting the critical importance of nucleolar activity during meiosis **(Figure 3, S4**). Collectively, these findings strongly suggest that a nucleolar stress response mechanism directly impairs meiotic progression, independently of downstream protein synthesis deficits.

### Chromatin remodeling drives meiotic defects in response to nucleolar disruption

Oogenesis requires elaborate chromatin remodeling, including gradual chromosome condensation during meiosis, and the spatial and temporal regulation of H3K4me3. Although many oogenic genes remain transcriptionally quiescent in late-stage oocytes, epigenetic priming at the chromatin level is essential for these genes to rapidly activate during zygotic genome activation. Thus, our study reveals a key non-ribosomal function of the RNA Pol I activity, and nucleolar integrity, in coordinating these critical epigenetic transitions.

Although nucleolar disruption clearly impacts chromatin organization, we did not observe a correlation between promoter accessibility and mature mRNA levels (**Figure S5**). This is consistent with another study in *C. elegans* that found no correlation between ATAC-seq signal and active transcription in isolated germ cells ^4^, suggesting that chromatin accessibility alone does not fully account for gene expression regulation in the germ line. Germline mRNA expression may instead be regulated by other mechanisms, including 22G-RNA and piRNA silencing (21U-RNAs) ^72–74^, inhibition of RNA Polymerase II carboxyl-terminal domain phosphorylation by PIE-1 ^75^, and post-transcriptional 3’UTR regulation controlled by RNA-binding proteins, such as PUF family proteins or GLD-1 ^76–79^. Indeed, prophase-arrested oocytes remain transcriptionally silent which coincides with PIE-1 re-localizing from the nucleolus to the nucleoplasm after nucleolar dissolution ^25^. Thus, premature nucleolar disruption may cause the untimely release of PIE-1, reinforcing transcriptional silencing despite increased chromatin accessibility. This mechanism likely explains why regions of differentially open chromatin do not become transcriptionally active upon RNA Pol I depletion.

Despite the lack of immediate transcriptional consequences, our data suggest that disruptions in chromatin conformation alone, driven by nucleolar cap formation, are sufficient to impair meiotic progression and reduce oocyte production. Given the extensive chromatin reorganization required in meiotic cells compared to other non-differentiating cell states, it is plausible that germ cells may be particularly vulnerable to such significant disturbances in nuclear architecture, even if these do not immediately affect transcriptional output.

### How nucleolar structure might drive meiotic chromatin dynamics

Two potential mechanisms may explain why nucleolar disruption impair meiotic progression, particularly through aberrant chromatin remodeling.

First, the disassembly of nucleolus via near complete depletion of RNA Pol I could release ribosomal proteins or other factors (e.g. MDM2) that, in other organisms, stabilize P53 or similar pathways to trigger cell cycle arrest ^80,81^. Although *C. elegans* lacks a direct MDM2 homolog, analogous pathways may be at play to prevent meiotic entry, potentially explaining the observed oogenesis defect. Indeed, our data show no significant stabilization of the C. elegans p53 ortholog, CEP-1, upon nucleolar disruption, indicating that any cell cycle arrest likely occurs independently of CEP-1/p53, possibly involving other checkpoint mechanisms or signaling pathways.

Second, nucleolar caps may function as stress sensors for DNA damage or incompletely replicated rDNA, activating surveillance pathways that stall meiotic progression. This model is supported by findings in other systems showing that nucleolar stress can prompt G2 arrest through ATR-Chk-1 signaling, independent of ribosome depletion ^82^. In *C. elegans*, rDNA loci may become exposed prematurely when nucleolar structure is compromised, leading to activation of ATM/ATR pathways and diverting double-strand break repair or chromatin-modifying complexes toward rDNA repeats. Indeed, we observed increased ATM/ATR checkpoint activation upon RNA Pol I depletion, supporting this interpretation. Given that proper ATM and ATR activity is critical for organization of chromosomes along the cohesin axis and normal meiotic progression ^83,84^, exposure of rDNA loci could disrupt the balance of ATM/ATR activity, leading to prophase arrest in RPOA-2-depleted animals.

An additional possibility is that aberrant chromatin remodeling, particularly premature deposition of H3K4me3 marks, could itself trigger ATM/ATR checkpoint signaling. Our results indicate that RNA Pol I depletion causes significant chromatin remodeling at promoters of oogenesis specific genes as well as other autosomal loci. This altered chromatin accessibility, driven by aberrant H3K4 methylation patterns, could disrupt normal synaptonemal complex formation and homologous recombination processes ^63^. Thus, nucleolar stress induced redistribution of chromatin modifying enzymes, including H3K4 methyltransferases, may initiate a cascade in which aberrant chromatin structure leads to synaptic defects and subsequent ATM/ATR-dependent checkpoint activation. However, since H3K4 methyltransferases can localize to double-strand break sites ^62^, this stress-driven redistribution of repair machinery might also underlie the misregulated H3K4me3 pattern observed in our study.

Ribosomal RNA transcription itself can also be regulated by epigenetic modifications at rDNA loci ^85^. Small molecule inhibitors such as CX-5461 or low dose oxaliplatin alter rDNA chromatin and activate DNA damage responses or senescence in a p53 independent manner ^86,87^. Since these drugs concurrently stabilize or damage G quadruplex structures and generate replication stress ^88,89^, their pleiotropic effects complicate interpretation. In contrast, our inducible genetic depletion of RNA Pol I subunits allows us to isolate the effects on genome-wide chromatin, revealing a clear link between Pol I loss, H3K4me3 mis-patterning, ATM/ATR activation and defective oogenesis, which occur independently of ribosome production.

### Spatiotemporal control of H3K4 methylation by the nucleolus during oogenesis

Our work demonstrates that premature nucleolar disassembly during oogenesis impairs germ cell maturation, primarily through its effects on chromatin remodeling. Specifically, the regulation of H3K4-related chromatin dynamics is critical for oogenesis, as H3K4me3 levels need to gradually increase along the distal-proximal axis to prime the promoters of genes required for zygotic genome activation ^4^. Disruptions in the activity of H3K4 de/methylases have been linked to abnormal nuclear morphology, reduced oocyte production, and inability to enter meiosis ^11–13^. In agreement, we find that premature nucleolar disruption weakens the H3K4 remodeling gradient, leading to defects in chromatin architecture and oocyte production. This highlights a novel role for nucleolar integrity as a critical upstream regulator of the spatiotemporal regulation of H3K4 methylation necessary for germ cell maturation.

Our gonadal ATAC-seq analyses further reveal that nucleolar disruption uniquely increases chromatin accessibility at oogenesis genes, oogenesis-promoting EFL-1-binding sites, and promoters undergoing H3K4me3 remodeling during oogenesis. We propose a model in which germ cells experiencing nucleolar stress redirect H3K4 methyltransferases or other remodeling factors to the site of rDNA damage, thereby depleting them from normal chromatin remodeling tasks. This shift may lead to hyper-accessible yet transcriptionally inactive loci in the germ line, consistent with the disruption of the normal H3K4me3 gradient and oogenesis-related gene priming.

Furthermore, our single-cell analysis of H3K4me3 immunofluorescence reveals that the positive H3K4me3 gradient along the distal-proximal gonad axis is disrupted under nucleolar stress. This perturbation is mediated by premature deposition of H3K4me3 in the distal region, further supporting the misregulation of chromatin accessibility in oogenesis genes. However, without single-cell ATAC-seq resolution, it remains unclear how chromatin accessibility changes are distributed along the gonad axis.

Although H3K4me3 represents a major dynamic mark in oocyte development, H3K27 methylation is also critical for germ line maintenance, and these marks are generally mutually exclusive ^4^. It remains possible that misregulated heterochromatin marks, such as H3K27me3, contribute to the nucleolar-disruption phenotype. Intriguingly, H3K27 methylation has been associated with the nucleolus in other contexts ^67^, further hinting that a global shift in histone marks underlies the oogenesis defects we observed.

### Broad implications

While our study focuses on *C. elegans,* nucleolar integrity and H3K4me3 remodeling during oogenesis are evolutionary conserved in higher eukaryotes, including humans ^5–10^. This suggests that nucleolus may play a similarly critical, non-ribosomal role in female gametogenesis in other species. The sensitivity of meiotic germ cells to nucleolar stress has significant implications for reproductive health ^90^, particularly in individuals exposed to environmental stress, undergoing aging, or affected by ribosomopathies.

Our findings also highlight potential explanations for phenotypic differences observed in disorders linked to Pol I components, such as Treacher Collins syndrome, which involves mutations in *POLR1B*, and uniquely impacts neural crest proliferation and migration ^91,92^. By uncovering how nucleolar driven chromatin reorganization shapes germ cell maturation, we provide a conceptual framework for understanding how disruptions in rDNA transcription can lead to specialized developmental deficits. More broadly, understanding these nucleolar roles offers a window into the intersection of genome maintenance, chromatin remodeling, and cell fate decisions across Metazoan development.

## ACKNOWLEDGEMENTS

We thank Trevor F. Freeman for generating ATAC-seq libraries. We thank Ayush Desai and Ajay Panda for their assistance with imaging experiments and vector construction used in this study, respectively. We thank Sarinay Cenik lab members, Cenik lab members, Can Cenik, Arlen Johnson, Steve Vokes, and Keiko Torii for constructive criticism and discussions.

This work was supported by the UT CNS Catalyst Grant, National Institutes of Health (NIH) NIGMS (R35GM138340), and Welch Foundation (F-2133-20230405) grants to E.S.C. Some strains were provided by the CGC, which is funded by NIH Office of Research Infrastructure Programs (P40 OD010440).

## AUTHOR CONTRIBUTIONS

R.M.T. and E.S.C. conceptualized the project and co-wrote the original manuscript. R.M.T., Q.Z., and E.S.C. revised the manuscript. Q.Z. and F.A. generated the RPOA-1 AID strain and dsRNAs. R.M.T. performed all imaging, immunostaining, data processing, and statistical analyses. Q.Z. performed crosses, injections, and all other molecular experiments. A.R. generated the ATAC-seq and RNA-seq libraries. E.S.C. supervised the project and acquired funding.

## RESOURCE AVAILABILITY

### Data availability

ATAC-seq and RNA-seq libraries are available through NCBI GEO, with accession codes GSE290498 and GSE290499, respectively.

### Code availability

Original pipeline used for germ nuclei linearization and confocal intensity quantification is hosted on the GitHub repository: https://github.com/raqmejtru/germ-nuclei-linearization.

## METHODS

### Strain generation

Constructs and strains used in this study are listed in **Table S7**-**S8**, respectively.

The *mKate2* tagged *nucl-1* allele was constructed using Cas9 protein driven by *eft-3* promoter in pDD162 and gRNA targeting a genomic sequence in the C-terminus of *nucl-1* in pAR2, a derivative of pRB1017, an empty vector for gRNA cloning. The sgRNA construct pAR2 was generated by the oligos ESC-AR-5 and ESC-AR-6. *nucl-1::mkate2::c1^sec^3xflag* (pAR1) was constructed for generating the knock-in into the C-terminus of the *nucl-1* gene. The 5′ and 3′ homology arms were amplified ∼500 bp upstream of *nucl-1* stop codon using oligos ESC-AR-7 and ESC-AR-8, and ∼500 bp downstream of stop codon using ESC-AKP-40 and ESC-AKP-41. The repair templates were used to replace the ccdB in pDD285.

Oligos ESC-AKP-(7-8) were used to generate a sgRNA targeting the N-terminus of the *dao-5* gene (16 bp from start codon) in pAKP2. The 5’ homologous arm (500 bp upstream of *dao-5* start codon) and 3’ homologous arm (500 bp downstream of start codon) were amplified using oligos ESC-AKP-(1-4), and subsequently used to replace the ccdB cassette in pDD282 to construct the repair template plasmid pAKP1.

The *degron*::*GFP*-tagged *rpoa-1* gene allele was constructed in a similar manner as above using pDD162 and gRNA targeting a genomic sequence in the C-terminus of *rpoa-1* in pQZ104. Oligos ESC-QZ-518 and ESC-QZ-519 were used to anneal sgRNA. The following reagents were used to assemble the final repair template pQZ103: 5′ homology arm (548 bp upstream of *rpoa-1* stop codon), 3′ homology arm (868 bp downstream of *rpoa-1* stop codon) were amplified using oligos ESC-QZ-512, ESC-QZ-513, ESC-FA-20, and ESC-FA-21. The 5′ homologue arm template was synthesized by a gBlock (IDT, ESC-QZ-511), while the 3′ homology arm was amplified using genomic DNA.

All plasmids for microinjection were purified using the PureLink HiPure Plasmid Miniprep Kit (Invitrogen #K210002). Oligo sequences used to generate these plasmids are in **Table S6**.

Strains generated through microinjection were prepared according to guidelines from ^93^. To generate ESC254, N2 animals were injected with a mix consisting of 50 ng/μL pDD162 (Cas9 vector), 50 ng/μL gRNA pAKP2, 50 ng/μL *dao-5* repair template of pAKP1, and 5 ng/μL of pCFJ104 as an extrachromosomal marker. The SEC was then excised by heat shock. To generate ESC770, N2 animals were injected with a mix consisting of 50 ng/μL pDD162, 50 ng/μL gRNA pAR2, 50 ng/μL *nucl-1* repair template of pAR1, and 5 ng/μL extrachromosomal marker L3785. Animals lacking the extrachromosomal array were heat-shocked to remove the SEC. To generate ESC772, DLW109 animals were injected with a mix consisting of 50 ng/μL pDD162, 50 ng/μL gRNA pRR13, 50 ng/μL *rpoa-2* repair template of pQZ43, and 5 ng/μL extrachromosomal marker pCFJ104. To generate ESC829, DLW109 animals were injected with a mix consisting of 50 ng/μL pDD162, 50 ng/μL gRNA pQZ104, 50 ng/μL *rpoa-1* repair template of pQZ103, and 5 ng/μL extrachromosomal marker pCFJ104. Finally, animals without the extrachromosomal array were heat-shocked to excise the SEC.

*WT/mIn1* males were crossed separately to ESC795, ESC772 and ESC796 strains to generate males carrying *EmGFP::syp-3* transgene and *mKate2*-tagged *nucl-1*, as well as hermaphrodites carrying *degron::GFP*-tagged *rpoa-2* or *degron::GFP*-tagged *grwd-1*. Both hermaphrodite genotypes were in the *BFP-*tagged *TIR1* and *WT/mIn1* background. These males were then crossed to the corresponding hermaphrodites to generate the ESC825 and ESC826 strains. We also crossed ESC772 to RB1025 to generate ESC818, a strain carrying *set-2(ok952)* mutation in auxin-inducible *rpoa-2* background. *set-2(ok952)* mutation was genotyped using the oligos ESC-QZ-515, 516, 517.

To visualize nucleolar structure while depleting RPOA-2 or GRWD-1, we crossed the *nucl-1::mKate2* reporter (ESC770) into the RPOA-2 AID background (ESC772) and GRWD-1 AID background (ESC796) to produce ESC794 and ESC797.

Lastly, to examine the co-localization of nucleolar structure and chromatin during meiosis, we crossed the *nucl-1::mKate2* reporter (ESC770) into a background containing a reporter for the synaptonemal complex (CA1218).

### Worm growth

*C. elegans* strains were maintained on plates with agar and nematode growth media (NGM) that were seeded with *Escherichia coli* strain OP-50. Animals were grown at 20°C for all experiments. Prior to experiments, animals were synchronized either by bleach synchronization or by allowing adults to lay embryos for two hours. The synchronized embryos were grown on NGM agar plates until they reached the late L4 stage. At the late L4 stage, animals were transferred to control or auxin treatment plates for further analysis.

### Auxin treatment

To deplete our proteins of interest using the auxin-inducible degron system (AID) system, we used a natural auxin (IAA) compound from Alfa Aesar (#A10556). A 400 mM stock solution was prepared in EtOH and stored at −20°C. For auxin treatment plates, we diluted IAA to a 1 mM concentration in NGM agar, then poured and let it solidify in plates. For control treatment plates, we diluted an equivalent volume of ethanol into NGM agar, then also poured and let it solidify in plates. Once solidified, plates were seeded with OP-50 and set to incubate at 20°C overnight to allow lawn growth. Synchronized late L4 worms were then transferred onto control or auxin plates for 18 hours at 20°C. Prior to any subsequent experiments, a qualitative assessment of GFP depletion was performed on a fluorescent dissection scope to ensure that auxin treatment worked as expected. Efficient degradation of RPOA-2 and GRWD-1 proteins were previously validated by western blotting within the experimental timeframe used in this study ^43,65^.

### Polysome fractionation

L4 larvae grown on regular NGM plates were transferred to NGM plates with and without 1 mM IAA for 24 hours at 20°C. Animals were liquid nitrogen flash frozen in polysome lysis buffer (20 mM Tris-HCl pH 7.4, 150 mM NaCl, 5 mM MgCl2) and ground in liquid nitrogen (with mortar and pestle). The frozen worm powder was thawed on ice and solubilized in polysome lysis buffer that was supplemented with 1 mM DTT, 100 μg/ml cycloheximide, 40 U/100 μl recombinant ribonuclease inhibitor (Invitrogen #10777019), 2 U/100 μl DNase (Invitrogen #AM2238). Lysates were loaded onto 10% to 45% sucrose gradients and spun for 2.5 hours at 40,000 rpm using SW 41 Ti rotor in an ultracentrifugation system (Beckman Coulter). RNA from monosome and polysome peaks was isolated using a density fractionation system (Brandel). The data were used for the analysis in **Figure S1E.**

The following steps were performed to calculate reductions in polysome abundance in auxin-treated samples relative to controls. We annotated thresholds on the fractionation curves generated above that encompassed free RNA regions, along with the first three polysome curves. Based on these thresholds, we calculated the area under the curve (AUC) for each region using the ‘AUC’ function in R with parameter ‘method = ‘trapezoid’’ ^94^. Normalized polysome abundance was calculated by dividing the polysome AUC by the free RNA AUC. We then compared this normalized abundance between auxin and control treatments.

### RNAi by dsRNA injection

We performed RNAi by injecting dsRNA into L4-stage animals. To generate the dsRNA, we amplified the following genes (*rpoa-2*, *grwd-1*, *fib-1*, and *nucl-1*) from cDNA, and non-target control (*wrmScarlet*) from plasmid pGLOW39. *T7* promoters were added to both ends of the amplified fragments using appropriate primers (ESC-RR-174/175, ESC-QZ-524/525, ESC526/527, ESC-QZ-528/529, ESC-QZ-416/417) (**Table S7**). The dsRNA synthesis was performed using the MEGAscript T7 Transcription Kit (Invitrogen #AMB13345) following the manufacturer’s instructions. We injected dsRNA (3µg/µl) into L4-stage animals of ESC772, COP262, TG12, and ESC795 strains. After injection, the animals were transferred to NGM plates and incubated at 20°C for 48 hours prior to imaging. At least 20 worms were injected for each dsRNA condition.

### Widefield and confocal imaging

Widefield and confocal images were captured using a STELLARIS 8 microscope (Leica Microsystems). When applicable, Z-stacks were transformed into average projections using LAS-X software (Leica Microsystems). For images that are presented for qualitative purposes, we adjusted the brightness and contrast to highlight features of interest, ensuring that adjustments are identical across treatments/conditions. Images used for statistical quantification were analyzed in their raw form without adjustments.

### Quantification of GFP signal

Imaging slides were prepared by transferring worms onto an immobilizing solution (10 mM levamisole hydrochloride, Sigma-Aldrich #31742-250MG). Gonads were dissected by using a hypodermic needle to slice near the head region. GFP signal was captured using a 10X objective with a GFP filter with consistent intensity and exposure settings.

CellProfiler v4.2.6 was used to segment individual nuclei and measure signal intensity based on the average signal of each segmented nucleus ^95^. Statistical comparison of mean GFP intensity between control and auxin treatments was carried out using a one-tailed Welch two-sample *t*-test, performed with the ‘t.test’ function in R ^94^.^87^

### Quantification of apoptotic corpses

Acridine orange (AO) staining was used to count apoptotic corpses in young adult germlines. First, L4s were transferred onto control or 1 mM IAA plates seeded with OP50 for 16 hours. We confirmed that the GFP signal was visibly reduced in the auxin-treated animals under a fluorescent dissection scope prior to continuing with AO staining. After 16 hours, each plate was spiked with 500 µL of AO stain prepared using 2 µL of 10 mg/mL AO (Invitrogen™ #A1301) dissolved in 1 mL of M9 buffer. Animals were left to feed on the stain for one hour, and were subsequently transferred onto new control or 1 mM IAA plates to de-stain for an additional hour before live imaging in levamisole.

Fluorescent microscopy was used to identify strong GFP signals characteristic of apoptotic corpses that retain a green AO signal. We counted the number of corpses per one gonad arm per animal, and used a Mann-Whitney *U*-test to compare the distribution of corpses across control and auxin-treated samples.

### Quantification of germline morphology

After control or auxin treatment, animals were transferred to slides with an immobilizing agent (10 mM levamisole hydrochloride, Sigma-Aldrich #31742-250MG). We captured whole-worm images for body measurements using a 10X DIC objective and germline morphology images using a 63X DIC objective. Using ImageJ, we measured body area, the length of the proximal arm (from the edge of the “-1” oocyte to the edge of the U-shaped arm bend), and the length and area of the three most proximal oocytes based on eggshell boundaries ^96^. Oocyte counts were measured by scanning vertical stacks for nuclear and eggshell boundaries while imaging. Statistical comparisons were performed with the ‘t.test’ function in R ^94^.

### Library preparation

After control or auxin treatment, worms were transferred onto a slide with an immobilizing solution (1 mM levamisole hydrochloride, Sigma-Aldrich #31742-250MG). Gonads were dissected by using a hypodermic needle to slice near the head region. Twenty dissected gonads were collected per replicate into ddH_2_O.

For RNA-seq, the dissected gonads were transferred to 1 mL of TRIzol (Thermo Fisher Scientific), vortexed, then incubated for 5 minutes at room temperature. RNA was extracted by adding 200 µL of chloroform to the lysate, spinning for 10 minutes at 15,000 rpm, and isolating the supernatant. The isolated RNA was precipitated overnight at −20°C using 50 mM sodium acetate, 5 mM MgCl_2_, 15 mg/mL GlycoBlue™ Coprecipitant (Thermo Fisher Scientific #AM9515), and isopropanol. RNA pellets were washed in 80% ethanol and prepared using the SMARTer Stranded RNA-Seq kit (Takara #634839). Briefly, the libraries were prepared by fragmenting and converting RNA to cDNA, and PCR amplified. PCR amplification reactions were run as follows: 1 minute denaturation at 94°C, followed by 15-17 cycles with 15 seconds at 98°C, 15 seconds at 55°C, and 30 seconds at 68°C. After PCR amplification, the DNA libraries were further purified using Agencourt AMPure XP beads (Fisher Scientific #A63880). The resulting libraries were quantified using the Qubit™ dsDNA HS Assay Kit (Thermo Fisher Scientific #Q32851) and sequenced on a NovaSeq 6000 v1.5 SP flow cell (Illumina).

For ATAC-seq, the dissected gonads were stained with DAPI (5 ng/µL in PBS), then loaded onto a hemocytometer for nuclei counting. A volume containing 50,000 nuclei was then mixed with 2.5 μL TDE1 tagmentase. The tagmentation reaction was incubated at 37 °C for 30 minutes on a shaker. Immediately after, the reactions were cleaned using the Zymo DNA Clean & Concentrator-5 kit (Fisher #NC9674845). The cleaned and tagmented DNA was then amplified with Nextera i5 and i7 adapters using the NEBNext® High-Fidelity 2X PCR Master Mix (NEB M0541L). Amplification reactions were run as follows: 5 minute extension at 72°C; 30 second denaturing at 98°C; 18-19 cycles with 10 seconds at 98°C, 30 seconds at 63°C, and 1 minute at 72°C; and a final extension of 10 minutes at 72°C. PCR products were cleaned to remove adapter contamination using the Zymo DNA Clean & Concentrator-5 kit. Library quality was assessed by checking nucleosome banding patterns on a gel, and passing libraries were subsequently cleaned using Agencourt AMPure XP beads. The resulting libraries were quantified using the Qubit™ dsDNA HS Assay Kit and sequenced on a NovaSeq 6000 v1.5 SP flow cell (Illumina).

### Quantification of rDNA transcription

Raw RNA-seq reads were trimmed with Trim Galore! v0.6.10 using default parameters ^97^. HISAT2 v2.2.1 was then used to build an index of the WBcel235 genome using the command ‘hisat2-build --seed 40’ ^98^. To report all multimapping alignments to repetitive rDNA regions, reads were mapped using the command ‘hisat2 --seed 40 --all’. Alignments were coordinate sorted using Picard SortSam v3.1.1 with the command ‘-SORT_ORDER coordinate --CREATE_INDEX truè ^99^.

To quantify rDNA transcription, we manually curated “gene” annotations for the two unique internal transcribed spacers (ITS) and appended them to the WBcel235 annotation. We assessed alignment counts to the modified annotation using featureCounts v2.0.6 with parameters ‘-t gene -g gene_id -M -O –fraction -p’ ^100^. A Welch two-sample *t*-test was performed to compare mean counts per million reads mapped to each ITS region as well as the entire 45S region using the ‘t.test’ function in R ^94^.

For a visual comparison of coverage in the rDNA region, we used bamCoverage v3.5.4 to normalize reads to average counts per million using the parameters ‘--normalizeUsing CPM --binSize 25’ ^101^. Coverage was later visualized in R using the Gviz v1.47.1 package ^102^.

rDNA transcription was also quantified by RT-qPCR. L4-stage animals were treated with or without auxin for 24 hours. Total RNA was treated with TURBO DNase (Thermo Fisher Scientific #AM2238), followed by phenol/chloroform extraction to remove DNA contamination. Approximately 800 ng of purified RNA was used for first-strand cDNA synthesis using SuperScript III Reverse Transcriptase (Thermo Fisher Scientific #18080-093) with random primers. RT-qPCR was performed using specific primer pairs (**Table S6**) on the QuantStudio 7 Flex Real-Time PCR System (Thermo Fisher Scientific). Relative rDNA transcript levels were determined by normalization to mature 26S rRNA.

### Identification of differentially expressed genes

Raw RNA-seq reads were trimmed with Trim Galore! v0.6.10 using default parameters ^97^. HISAT2 v2.2.1 was then used to build an index of the WBcel235 genome using the command ‘hisat2-build --seed 40’ ^98^. Reads were mapped using the command ‘hisat2 --seed 40’ with default parameters. Alignments were coordinate sorted using Picard SortSam v3.1.1 with the command ‘-SORT_ORDER coordinate -- CREATE_INDEX truè ^99^. Alignment counts were then summarized using featureCounts v2.0.6 with parameters ‘-t exon -g gene_id -p’ ^100^. Before further analyses, we filtered genes down to the 19,985 protein coding genes found in the WBcel235 annotation, and we also filtered out genes with low expression (less than 5 counts across 3 biological replicates). We evaluated differential gene expression using DESeq2 v1.40.2 with an experimental design that accounted for strain and treatment ^103^. Genes were considered differentially expressed if they had a Benjamini-Hochberg adjusted *p*-value < 0.05.

### Identification of differentially accessible regions

Raw ATAC-Seq reads were trimmed, aligned, decontaminated, and quality assessed using the SRAtac v0.5.0 pipeline ^65^. Before performing differential accessibility analyses, we used RUVSeq v1.34.0 to perform batch correction on mapped counts by computing estimated factors of unwanted variance against an empirical set of control genes ^104^. We arbitrarily selected control genes with *p*-values greater than 0.75 and an average count per million above 20 in at least 90% of samples, calculated with the ‘filterByExpr’ function in edgeR v3.42.4 ^105^.

To evaluate differentially accessible chromatin regions (DARs), we used DESeq2 v1.40.2 with an experimental design that accounted for strain, treatment, and the estimated factors of unwanted variation ^103^. We compared DARs across non-depleted and depleted samples using a significance threshold of a Benjamini-Hochberg adjusted *p*-value < 0.05. To visualize DARs, we performed log_2_ fold-change shrinkage using DESeq2’s ‘lfcShrink’ function with apeglm v1.22.1 shrinkage estimators ^106^.

### ATAC-seq genomic feature annotation

ATAC-seq peaks were annotated using the ‘annotatePeak’ function in the ChIPseeker v1.36.0 package with modification ‘overlap = all’ ^107^ in conjunction with the TxDb ce11 v3.4.6 reference database ^108^. Results were visualized with the ‘plotAnnoBar’ function in the clusterProfiler v4.12.0 package ^109,110^.

### Differential analysis of functional gene ontology (GO) biological processes

For both ATAC-seq and RNA-seq, differential analysis of functional gene ontology (GO) biological processes were performed using the ‘compareCluster’ function in the clusterProfiler v4.12.0 package using the following parameters: fun = ‘enrichGO’, ont = ‘BP’, pAdjustMethod = ‘fdr’, pvalueCutoff = 0.0001 (ATAC) or 0.001 (RNA), qvalueCutoff = 0.0001 (ATAC) or 0.001 (RNA) ^109,110^. Results were visualized using the ‘dotplot’ function in the enrichplot v1.24.0 package using parameter showCategory = 3 ^111^.

### Motif enrichment and characterization

The MEME-suite bed2fasta tool was used to add nucleic sequences to background and differentially accessible peaks based on the UCSC ce11 reference genome ^112^. We used the MEME-suite STREME tool to enrich for ungapped motifs present in our peak sets with a *p*-value and *E*-value threshold of 0.05, an alignment around the center of the peak, and all other default parameters ^113^. We used the MEME-suite Tomtom tool ^114^ to map the enriched motifs against the 842 TF motifs present in the TFBSshape database ^54^. We classified significant motif-motif similarity based on Pearson correlation coefficients, an *E*-value threshold of 8.42, and all other default parameters. We further manually curated matches to combine redundant query motifs and their repeated hits, and to ensure that alignments were visibly cohesive.

To identify genomic sites in our ATAC-seq data containing the *C. elegans* EFL-1 motif (MA0541.1), we used the MEME-suite FIMO v5.5.5 tool with parameters ‘--thresh 1.0E-4 --nrdb--’ against our background set of peaks (i.e., all peaks across all samples called in SRAtac pipeline). We identified which peaks contained EFL-1 motifs by searching for overlaps between EFL-1 motif sites and peaks using the ‘findOverlaps’ function with parameter ‘type = ‘within’’ in the GenomicRanges v1.56.1 package ^115^. Genes with EFL-1 motifs in their promoter were analyzed for enrichment of GO biological processes using the enrichGO function in the in the clusterProfiler v4.12.0 package with parameters ‘pAdjustMethod = ‘fdr’, pvalueCutoff = 0.05, qvalueCutoff = 0.05’ based on the annotation of the peak that the motif was located in ^109,110^. Results were visualized using the ‘dotplot’ function in the enrichplot v1.24.0 package using parameter showCategory = 3 ^111^.

### Metagene analysis of histone ChIP-seq coverage

Isolated germ and oocyte histone ChIP-seq experiments used in this study are listed in **Table S9**.

Raw ChIP-seq reads were downloaded from NCBI using the SRA Toolkit ^116^. Reads were processed using the nf-core/chipseq pipeline with parameters: ‘--genome WBcel235 --broad_cutoff 0.01 --macs_fdr 0.001 --nomodel’ and ‘--extsizè between 147-250 ^117,118^. If read length was known, we specified the ‘--read_length’ parameter, otherwise we specified the effective genome size (‘--macs_gsize 100286401’) estimated using the recommended ‘faCount’ tool ^119^.

Next, bigwigAverage v3.5.5 was used to compute an average bigwig coverage track for each antigen and cell type combination (i.e., oocyte H3K4me3 antigen, oocyte H3K4me3 input, etc.) using parameter ‘--scaleFactors’ with the scale factors provided by the nf-core/chipseq pipeline ^101^. Average log_2_ ratios of coverage across antigen/input samples were then computed using bigwigCompare v3.5.5 with parameter ‘--operation log2’ ^101^. Additionally, an average log_2_ coverage score was computed across significantly more accessible peaks after RPOA-2 depletion and an identically-sized random subset of ATAC-seq peaks by intersecting bigwig scores using the ‘findOverlaps’ function in the GenomicRanges v1.56.1 ^115^.

Metagene profile plots were used to display the average log_2_ ratios of ChIP signal from isolated germ cells or oocytes across peaks that were significantly more accessible after RPOA-2 depletion, or across an identically-sized random subset of promoter ATAC-seq peaks that were not significantly more accessible. The tool computeMatrix v3.5.5 was used to generate coverage scores per peak using parameters ‘-- referencePoint center --afterRegionStartLength 1000 --beforeRegionStartLength 1000 --binSize 25’ ^101^. Metagene plots were displayed using plotProfile v3.5.5 with parameter ‘--perGroup’ to summarize scores across germ versus oocyte cell types ^101^.

### Analysis of chromatin accessibility across H3K4me3-remodeled gene clusters

Scaling (size) factors for ATAC-seq peak counts were computed using the relative log expression (RLE) normalization method from DESeq2 v1.40.2 ^103^. Each sample’s ATAC-seq coverage was re-scaled and binned into bigwigs for downstream visualization using bamCompare v3.5.5 with parameters ‘--scaleFactor (1/RLE size factor) --binSize 50’ ^101^. RLE-normalized coverage for depletion and control samples was aggregated using bigwigAverage v3.5.5, and subsequently log_2_ transformed using bigwigCompare v3.5.5 ‘--operation log2’ to interpret fold changes in coverage across depletion and control conditions ^101^.

Metagene profile plots were used to display the average log_2_ ratios of ATAC signal across the H3K4me3-remodeled gene clusters. The tool computeMatrix v3.5.5 was used to generate coverage scores per gene using parameters ‘scale-regions --afterRegionStartLength 1000 --beforeRegionStartLength 1000 -- regionBodyLength 1000 --binSize 25’ ^101^. Metagene plots were displayed using plotProfile v3.5.5 with parameter ‘--perGroup’ to summarize scores across cluster 1, 2, and 3 gene sets ^101^.

### Gonadal immunostaining

Prior to immunostaining, we confirmed that degron::GFP::RPOA-2 signal was visibly reduced in auxin-treated animals.

H3K4me3 immunostaining was performed on tube-fixed dissected gonads from adult worms after control or auxin treatment. Briefly, gonads were dissected in a solution with levamisole and 0.05% Tween 20 in PBS (PBS-T). Dissected bodies were transferred to DNA LoBind™ Tubes (Eppendorf™ #022431021) and fixed for five minutes using 4% PFA, then washed three times with PBS-T. Next, gonads were permeabilized for fifteen minutes using 0.5% Triton X-100 in PBS-T. Gonads were then post-fixed using 95% ethanol for five minutes at -20°C, washed three times in PBS-T, and blocked for one hour using 5% BSA in PBS-T. For primary antibody incubation, half of the gonads from each treatment condition were incubated overnight at 4°C in 2% BSA in PBS-T as a negative primary antibody control, while the other half were incubated overnight at 4°C using the primary antibody (Rabbit Anti-Histone H3 (tri-methyl K4) antibody (Abcam #ab8580), 1:250; Rabbit Phospho-(Ser/Thr) ATM/ATR Substrate Antibody (Cell Signaling #2851), 1:250) in 2% BSA in PBS-T. Gonads were subsequently washed three times using PBS-T and incubated at room temperature for two hours in the secondary antibody (For H3K4me3: 10 μg/mL of Goat Anti-Rabbit IgG (H+L) Cross-Absorbed Alexa Fluor™ 633 (Invitrogen™ #A-21070); for Phospho-(Ser/Thr) ATM/ATR Substrate: Goat anti-Rabbit IgG (H+L) Cross-Adsorbed Secondary Antibody, Alexa Fluor™ 555 (Thermo Fisher Scientific #A-21428), 1:200) in 2% BSA in PBS-T. Finally, gonads were washed three times with PBS-T and mounted onto slides using VECTASHIELD® Antifade Mounting Medium with DAPI (1.5 μg/mL) (Vector Laboratories #H120010).

Immunostained gonads were imaged from the distal to proximal zones using 40X tile-scanned confocal Z-stacks of identical resolutions (2048x2048). All Z-stacks were forced to encompass 3 μm to minimize overlaps between nuclei for downstream nuclei segmentation. Additionally, all confocal images were captured using identical laser intensity and gain settings for downstream intensity quantification. We excluded imaging gonads that were not adequately laying flat on the slide, preventing us from completely capturing the distal to proximal zones within a 3 μm vertical space. Lastly, Z-stacks were transformed into average projections for further analyses using LAS-X software (Leica Microsystems).

### Gonad linearization and H3K4me3 signal quantification

A custom QuPath v0.5.1 script was created to identify H3K4me3 signal intensity per nuclei following RPOA-2 control or depletion treatment. Briefly, gonads were manually annotated based on morphology from the T-PMT confocal channel, and these annotations were overlaid onto the corresponding DAPI channel. Gonad annotations were segmented using the ‘StarDist2D’ function on the DAPI channel with the ‘dsb2018_heavy_augment.pb’ model and parameters ‘normalizePercentiles(1, 99)’, ‘threshold(0.80)’, ‘pixelSize(0.1)’, and ‘cellExpansion(0)’ ^120,121^. Segmented nuclei annotations were copied to the H3K4me3 channel. Average H3K4me3 and DAPI signal intensities within segmented nuclei were measured using the ‘addIntensityMeasurements’ function with parameter ‘downsample = 0.1’. We converted average signal intensity to integrated signal intensity using the formula ‘integrated intensity = mean intensity * area of detected nucleì to prevent nuclei detection area from biasing intensity comparisons across differently sized nuclei. Additionally, to account for differences in permeability across gonads during immunostaining (due to varying physical damage while vortexing, etc.), we normalized integrated H3K4me3 intensity to integrated DAPI signal, and separately to the maximum H3K4me3 intensity recorded per gonad.

To analyze variation in H3K4me3 intensity from the distal to proximal zones, we employed a variation of a gonad linearization algorithm to track the location of each nuclei along a linearized gonad representation ^122^. Briefly, in QuPath, a polyline skeleton was drawn from the distal tip to the most proximal end of the gonad. We computed perpendicular projections of each nucleus onto the nearest gonad skeleton segment and used this value to calculate the position of each nucleus along the gonad skeleton. We limited our analyses to nuclei that were within a linearized gonad length of 2,000 px.

Lastly, for each gonad, we evaluated the positive non-parametric correlation between nuclei position along the linear gonad axis and integrated H3K4me3/DAPI intensity or per-gonad scaled H3K4me3 intensity using the ‘cor.test’ function in R with parameters ‘method=’spearman’, alternative=’greater’, exact=’false’’ and corrected for false positives using the ‘p.adjust’ function in R with parameter ‘method=’BH’’ ^94^.

## SUPPLEMENTAL TABLE LEGENDS

Table S1. Differential accessibility results from DESeq2 analysis comparing RPOA-2 control vs depletion conditions, as well as GRWD-1 control vs depletion conditions.

Table S2. Gene ontology (GO) biological processes enriched in gene sets with increased accessibility after reducing RNA Pol I activity or ribosome biogenesis. (GeneRatio: the proportion of annotated genes that are significantly more accessible and belong to a specific GO set, relative to the total number of significantly more accessible genes. BgRatio: the proportion of genes in a specific GO set, relative to the total number of annotated *C. elegans* genes).

Table S3. Ungapped sequence motifs enriched in differentially accessible regions (DAR) after reducing RNA Pol I activity or ribosome biogenesis, compared to motifs in background peaks. (Sites: the number and percentage of ATAC-seq regions in a DAR set that match the motif). (S.M.A. = significantly more accessible peaks, S.L.A. = significantly less accessible peaks)

Table S4. Differentially expression results from DESeq2 analysis comparing RPOA-2 control vs depletion conditions, as well as GRWD-1 control vs depletion conditions.

Table S5. Gene ontology (GO) biological processes enriched in differentially expressed genes (DEG) after reducing RNA Pol I activity or ribosome biogenesis. (GeneRatio: the proportion of DEGs that belong to a specific GO set, relative to the total number of DEGs. BgRatio: the proportion of genes in a specific GO set, relative to the total number of annotated C. elegans genes).

Table S6. Oligonucleotides used in this study.

Table S7. Constructs used in this study.

Table S8. Strains generated and used in this study. Table S9. Data sets used in this study.

---

**Figure S1.**
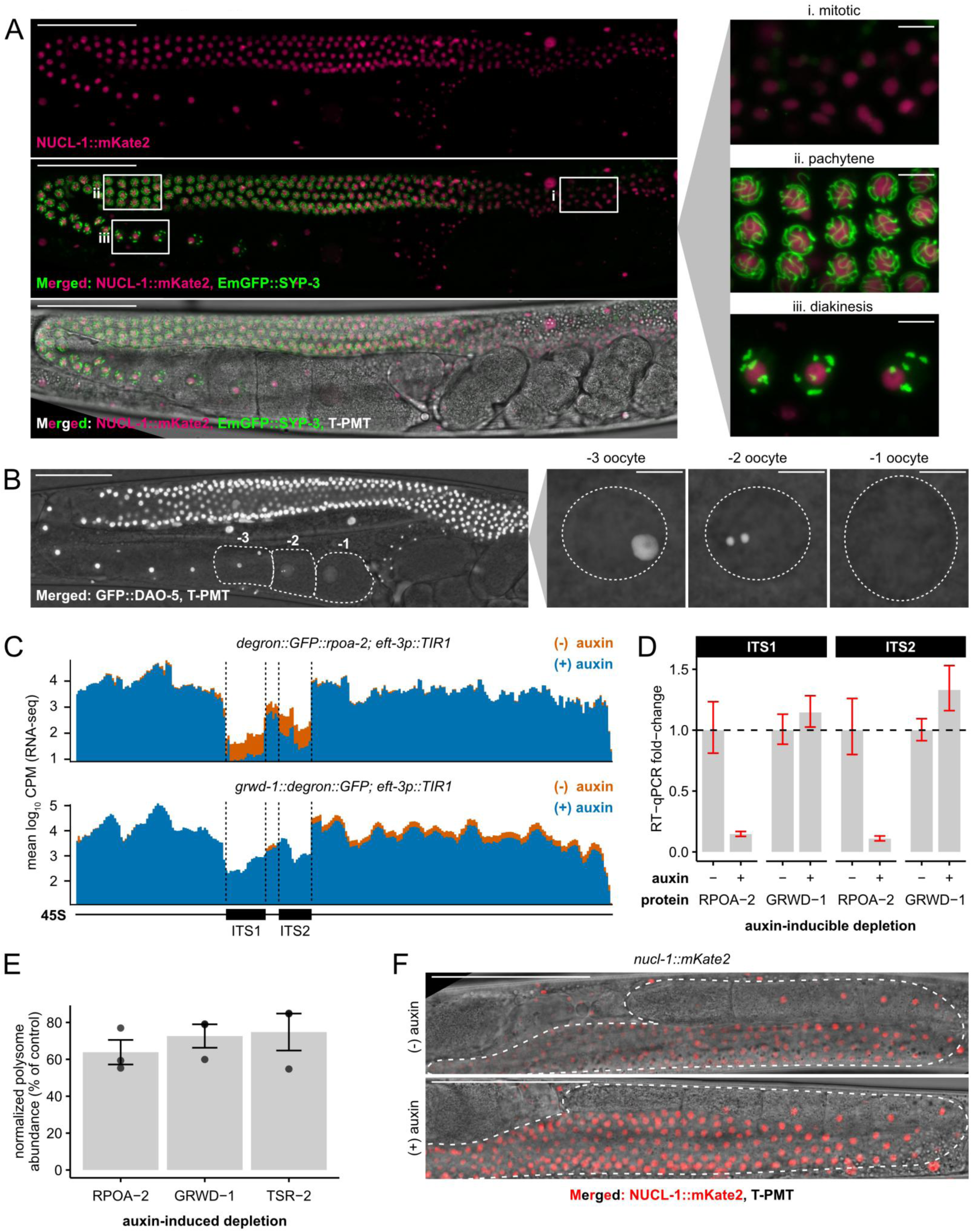
**Related to Figure 1. Nucleolar dynamics throughout the germ line (A, B, F); effects of Pol I and ribosome assembly chaperone depletion on the transcription of internally transcribed spacers (ITS) and downstream polysome abundance (C-E).** (**A**) Average confocal projection of an adult germ line (left) with an endogenous NUCL-1::mKate2 nucleolar marker and an EmGFP::SYP-3 meiosis marker. Nucleoli are present throughout the mitotic, pachytene, and diakinesis zones (right), as indicated by the localization of EmGFP::SYP-3. Scale bar = 50 μm (full germ line) and 5 μm (zoom). (**B**) Average confocal projection of an adult germ line (left) with an endogenous GFP::DAO-5 nucleolar marker. Zoomed in nuclei (right) reveal that the nucleolus forms droplets in the “-2” oocyte and is absent in the most proximal and mature “-1” oocyte. Scale bar = 50 μm (germ line) and 5 μm (nuclei). (**C**) Average log_10_ counts per million (CPM) reads from gonadal RNA-seq (total RNA) mapped to a simplified single-copy 45S rDNA locus. Averages are composed of three biological replicates per condition, each consisting of 20 gonads. RPOA-2-depleted germ lines show reduced coverage across internally transcribed spacer (ITS) regions compared to GRWD-1-depleted germ lines. (**D**) RT-qPCR analysis of whole-animal ITS1 and ITS2 expression following auxin-inducible depletion of RPOA-2/GRWD-1. Relative transcript levels of ITS1 and ITS2 were normalized to mature 26S rRNA. Data represent three biological replicates for RPOA-2 depletion and two biological replicates for GRWD-1 depletion, with three technical replicates per biological replicate. Bars and error bars represent mean ± SD. (**E**) Polysome abundance after auxin-induced depletion of RPOA-2, GRWD-1, and TSR-2, normalized to the abundance of free RNA. Points represent biological replicates (3 per condition); bars and error bars represent the mean and SEM, respectively. (**F**) NUCL-1::mKate2 is morphologically unaffected by 1 mM IAA in a wild-type background. Scale bar = 50 μm.

**Figure S2.**
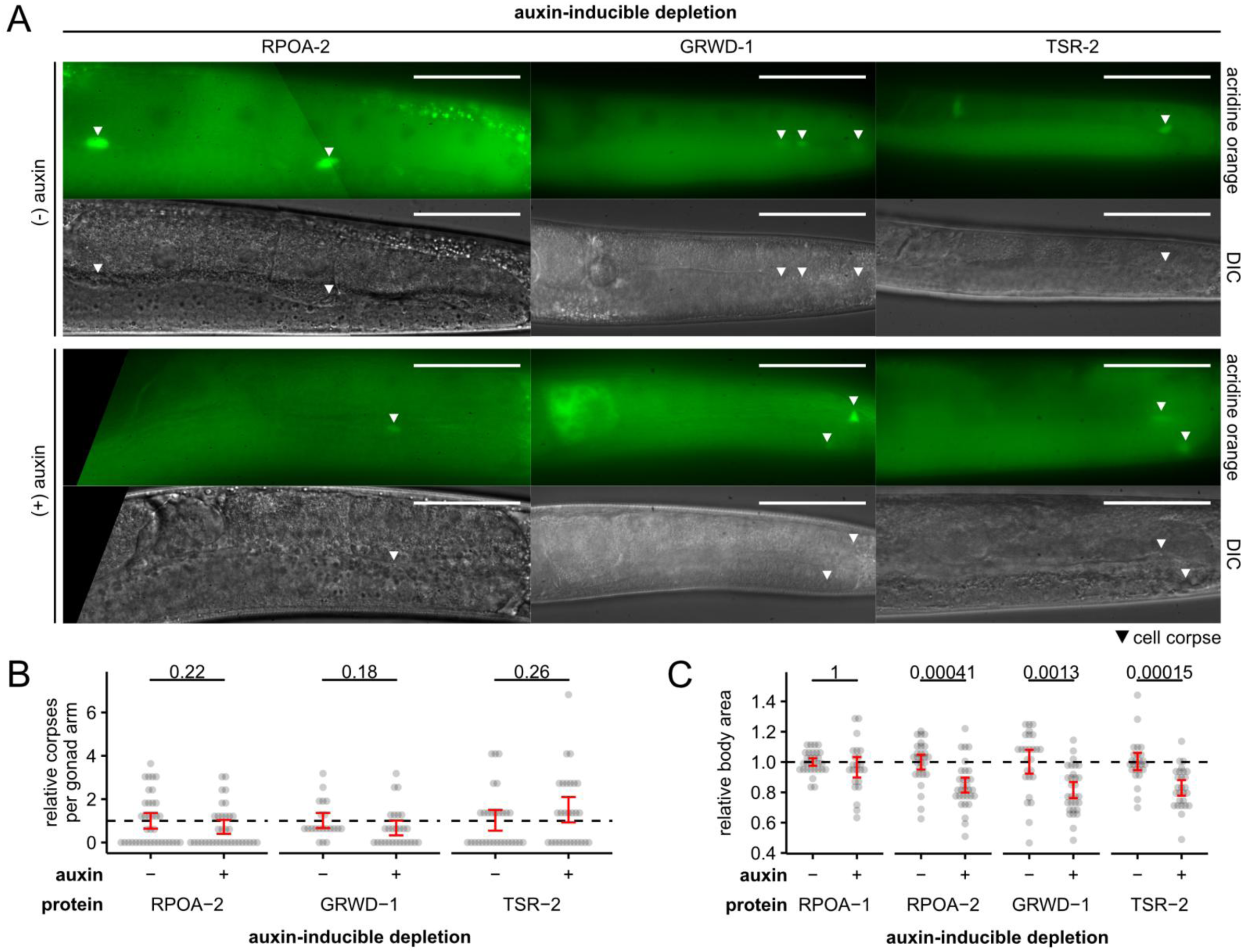
**Related to Figure 2. Effects on germ cell death and animal body area under reduced RNA Pol I activity or ribosome assembly.** (**A**) Acridine-orange-stained germ lines after auxin-inducible depletion of RPOA-2/GRWD-1/TSR-2. Cell corpses are indicated by white triangles; gonad arms are outlined in white. Scale bar = 50 μm. (**B**) Distribution of cell corpses per gonad arm based on acridine orange staining, normalized to the mean of each strain’s control group. Each point represents the number of corpses in a gonad arm across three biological replicates. Statistical comparisons performed using a two-tailed Welch’s two-sample *t*-test. (**C**) Distribution of body areas after auxin-inducible depletion of RPOA-1/RPOA-2/GRWD-1/TSR-2. Each point represents measurements from an individual worm, normalized to the mean of each strain’s control group. Crossbars represent SEM. Statistical comparisons performed using a left-tailed Welch’s two-sample t-test with Bonferroni corrections.

**Figure S3.**
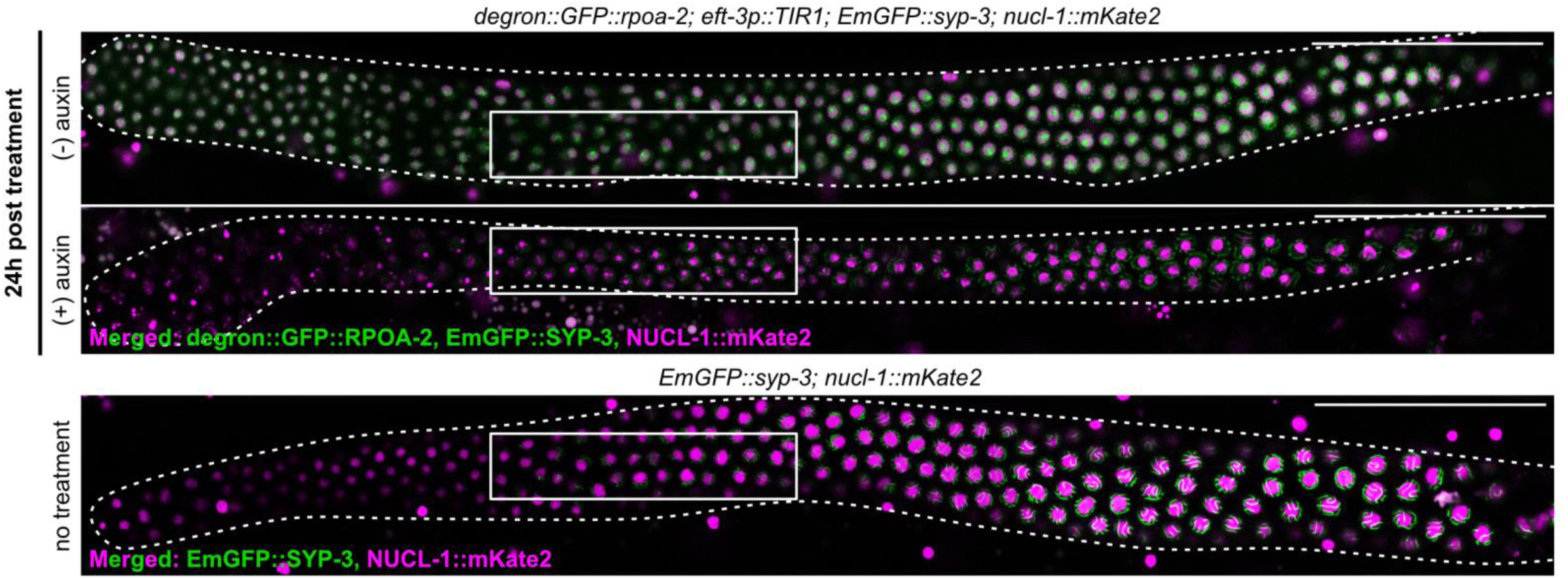
**Related to Figure 3. Synaptonemal complex formation under reduced Pol I activity during the mitotic-to-meiotic transition in an adult germ line.** Confocal images of adult distal germline arms (dotted lines) expressing degron::GFP::RPOA-2, EmGFP::SYP-3, and NUCL-1::mKate2 following control or auxin-inducible depletion of RPOA-2 (top rows). Additionally, we show a non-treated control in a wild-type genetic background (bottom row). White rectangles show zoomed-in transition zone regions in Figure 3A. Scale bar = 50 μm.

**Figure S4.**
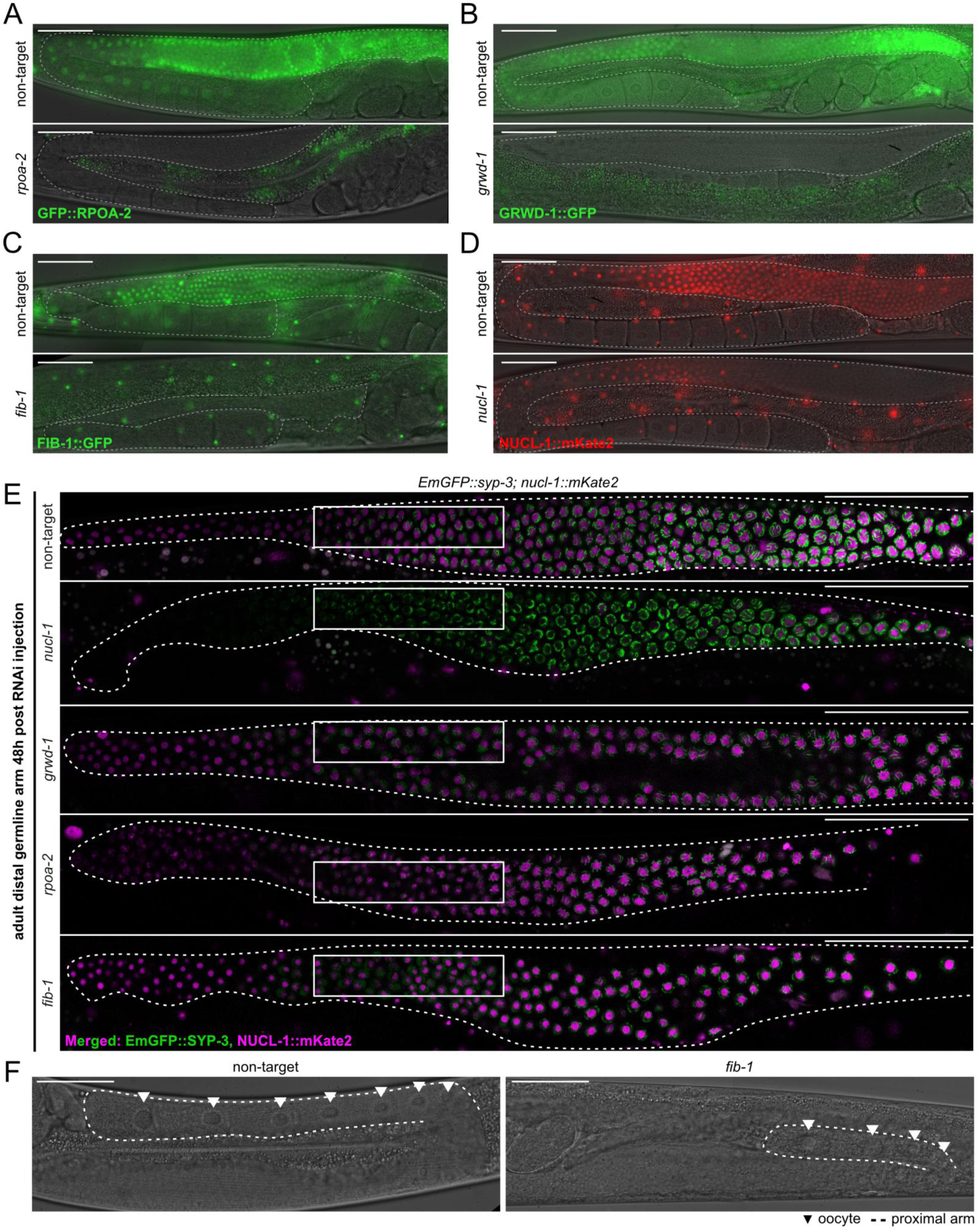
**Related to Figure 3.** Examining germline synapsis and proximal oocytes after knockdown of nucleolar proteins. **(A-D)** Merged fluorescent and DIC images of germ lines (outlined in white) after RNAi injection targeting *rpoa-2* (A), *grwd-1* (B), *fib-1* (C), and *nucl-1* (D), as well as non-target (*wrmScarlet*) injection controls. Each of the endogenous fluorescent-tagged proteins of the RNAi target genes show reduced germline expression after 48 hours. Scale bar = 50 μm. **(E)** Confocal images of adult distal germline arms expressing EmGFP::SYP-3 and NUCL-1::mKate2 48 hours post RNAi knockdown of a non-target locus (*wrmScarlet*), *nucl-1*, *grwd-1*, *rpoa-2*, and *fib-1*. White rectangles show zoomed-in transition zone regions in Figure 3B. Scale bar = 50 μm. All images were taken with identical laser intensity; due to decreased SYP-3 expression under knockdown of *grwd-1, rpoa-2,* and *fib-1*, we have increased the brightness of these conditions to be more comparable to wild type. **(F)** DIC images of adult proximal germline arms (dotted line) and proximal oocytes (triangles) 48 hours post RNAi knockdown of a non-target locus (*wrmScarlet*) and *fib-1*. Scale bar = 50 μm.

**Figure S5.**
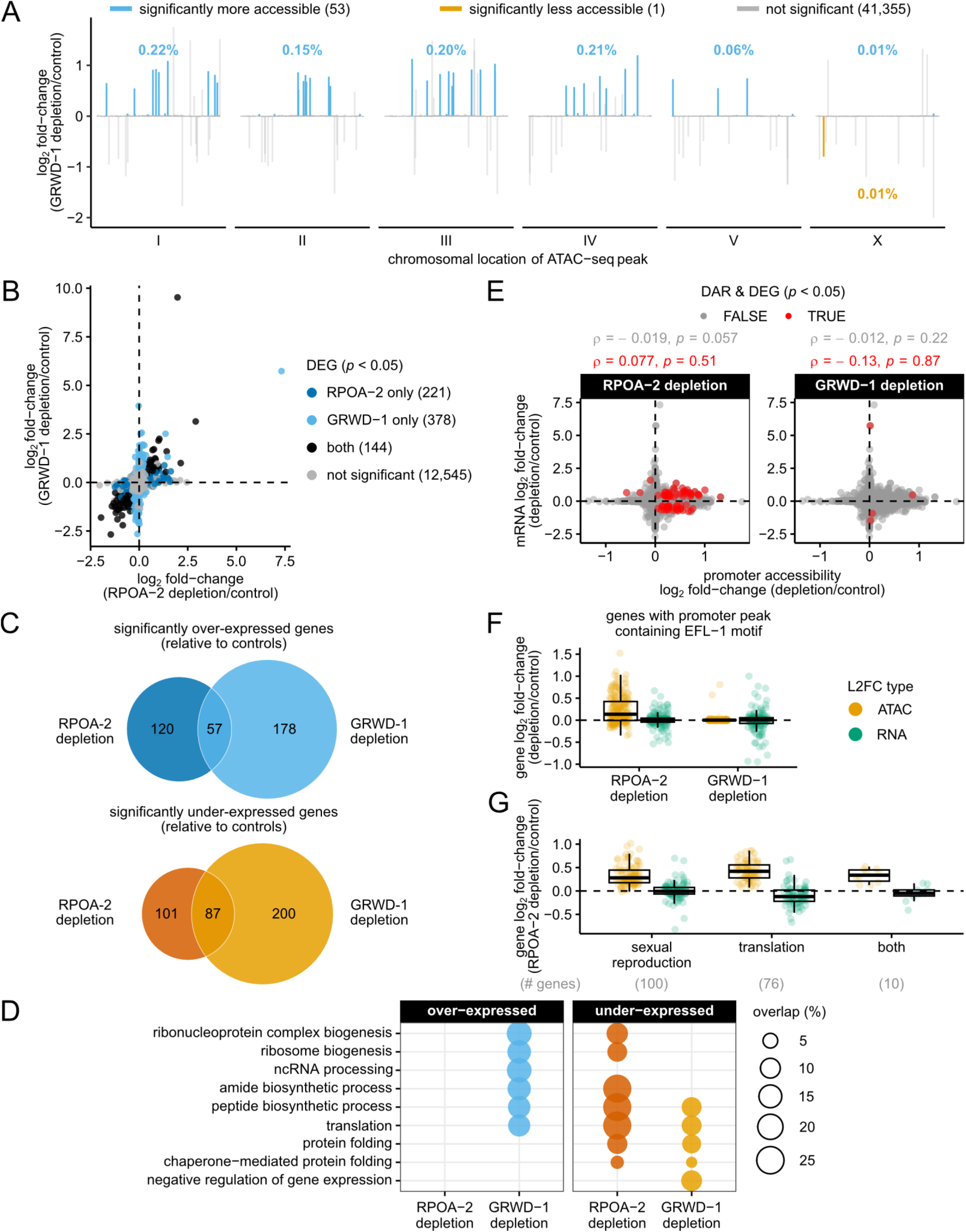
**Related to Figure 4. Effects of RPOA-2 and GRWD-1 depletion on germline chromatin accessibility and gene expression.** (**A**) Log_2_ fold-change estimates of chromatin accessibility based on gonadal ATAC-seq data from at least three biological replicates per condition, each composed of 20 gonads. Each segment along the x-axis represents a genomic region. The number of differentially accessible regions are noted in the plot legend based on an adjusted *p* < 0.05. Percentages represent the proportion of differentially accessible peaks relative to the total number of called peaks per chromosome. (**B**) Log_2_ fold-change estimates of gene expression from RNA-seq conducted on dissected gonads after RPOA-2 or GRWD-1 depletion. Biological replicates, consisting of 20 gonads: RPOA-2 control = 3, RPOA-2 depletion = 4, GRWD-1 control = 2, GRWD-1 depletion = 3. (**C**) Venn diagram to contextualize the difference in the scale of gene expression changes after RPOA-2 depletion compared to GRWD-1 depletion. (**D**) Enrichment of gene ontology biological processes in genes that were differentially expressed after RPOA-2 or GRWD-1 depletion (Hypergeometric test, FDR < 0.001). (**E**) Log_2_ fold-change estimates of promoter accessibility versus mRNA to investigate whether chromatin accessibility correlated with gene expression. We did not identify a significant correlation between promoter accessibility and mRNA levels across all genes (gray), or across genes that were both differentially accessible and differentially expressed after RPOA-2 or GRWD-1 depletion (black). (**F**) Comparison of ATAC-seq and RNA-seq log_2_ fold-change estimates for genes with promoter peaks containing an EFL-1 motif after RPOA-2 or GRWD-1 depletion. On average, genes with a promoter EFL-1 motif did not show a significant increase in mRNA fold-changes after RPOA-2 depletion (one-tailed one-sample *t*-test, alternative hypothesis: log_2_ fold-change > 0, *p* = 0.14). (**G**) Comparison of ATAC-seq and RNA-seq log_2_ fold-change estimates in biological processes enriched in genes with increased chromatin accessibility after RPOA-2 depletion. Only the terms “sexual reproduction” and “translation” are represented, as they encompass the majority of related subprocesses. “Both” represents genes that are involved in both biological processes. On average, translation-related genes, including ribosomal proteins, have lower mRNA fold-changes after RPOA-2 depletion (one-tailed one-sample *t-*test, alternative hypothesis: average log_2_ fold-change < 0, *p* = 0.00020), whereas genes related to sexual reproduction do not have significantly increased mRNA levels (one-tailed one-sample *t*-test, alternative hypothesis: log_2_ fold-change > 0, *p* = 0.16).

**Figure S6.**
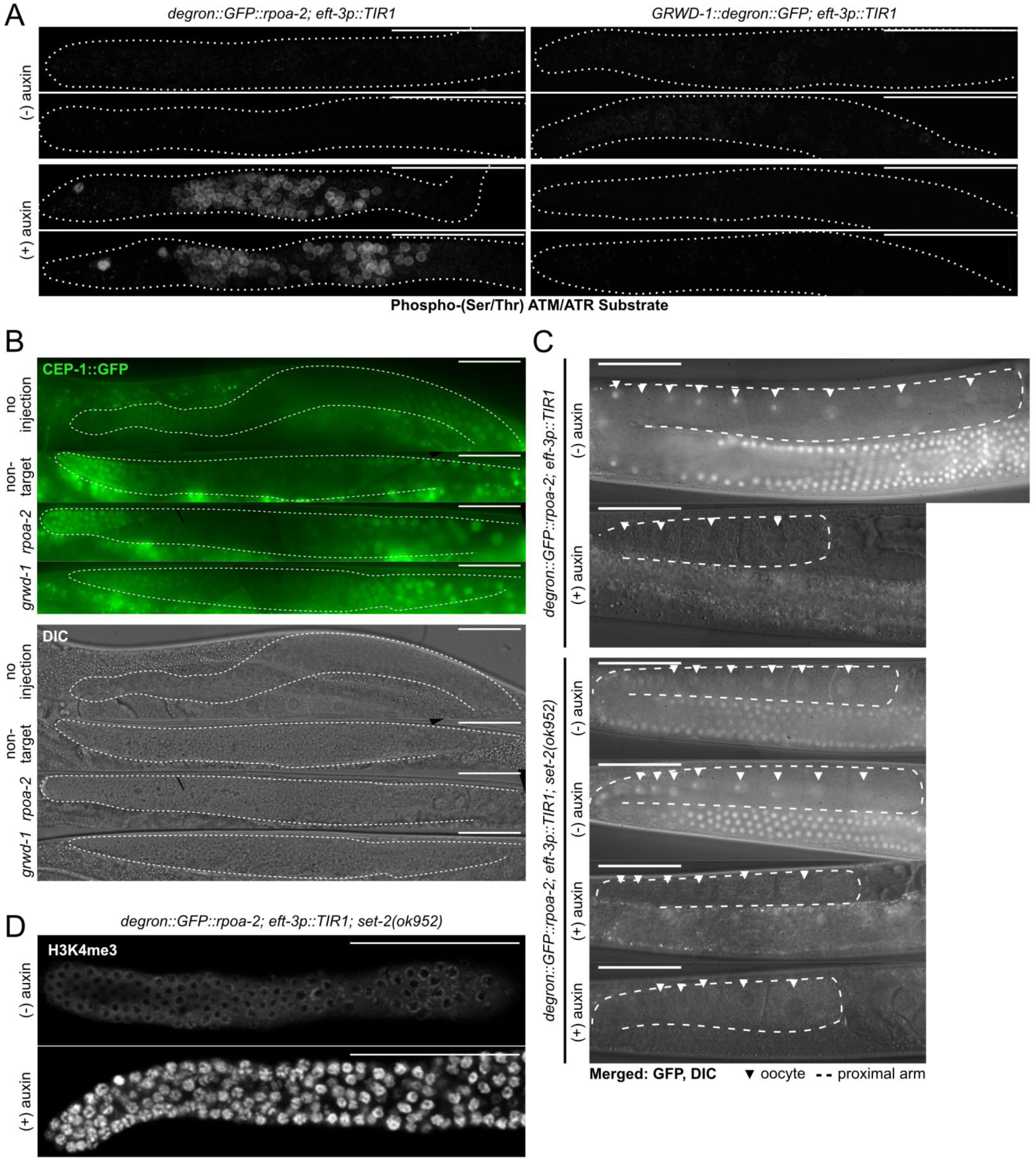
**Related to Figure 5. Pol I reduction uniquely triggers ATM/ATR signaling in a CEP-1 independent manner (A-B); H3K4 trimethylation increases under reduced Pol I activity in a SET-2 independent manner (C-D).** (**A**) Average confocal projections of dissected gonad arms (outlined in white) immunostained for Phospho-(Ser/Thr) ATM/ATR Substrate after auxin-inducible depletion of RPOA-2/GRWD-1. All conditions were imaged with identical laser settings. Scale bar = 50 μm. (**B**) Fluorescent and DIC images of CEP-1::GFP expression in the germ line (outlined in white) after RNAi injection targeting *rpoa-2* and *grwd-1*, as well as non-target injection and no injection controls. All conditions imaged with identical settings. Scale bar = 50 μm. (**C**) Merged fluorescent (degron::GFP::RPOA-2) and DIC images of adult germ lines after auxin-inducible depletion of RPOA-2 in a wild-type or *set-2(ok952)* genetic background. Dotted lines highlight the proximal arm; triangles point to proximal oocytes. Scale bar = 50 μm. (**D**) Representative average confocal projections of adult gonad arms that have been immunostained for H3K4me3 after auxin-inducible depletion of RPOA-2 in a *set-2(ok952)* genetic background. Dotted lines represent gonad boundaries determined from T-PMT. Scale bar = 50 μm.

## REFERENCES

1. Hamatani T, Carter MG, Sharov AA, Ko MSH. Dynamics of Global Gene Expression Changes during Mouse Preimplantation Development. Developmental Cell. 2004;6(1):117–131. doi:10.1016/S1534-5807(03)00373-3

2. Vastenhouw NL, Cao WX, Lipshitz HD. The maternal-to-zygotic transition revisited. Development. 2019;146(11):dev161471. doi:10.1242/dev.161471

3. Tadros W, Lipshitz HD. The maternal-to-zygotic transition: a play in two acts. Development. 2009;136(18):3033–3042. doi:10.1242/dev.033183

4. Mazzetto M, Gonzalez LE, Sanchez N, Reinke V. Characterization of the distribution and dynamics of chromatin states in the C. elegans germline reveals substantial H3K4me3 remodeling during oogenesis. Genome Res. Published online December 26, 2023. doi:10.1101/gr.278247.123

5. Blythe SA, Cha SW, Tadjuidje E, Heasman J, Klein PS. β-catenin Primes Organizer Gene Expression By Recruiting a Histone H3 Arginine 8 Methyltransferase, Prmt2. Dev Cell. 2010;19(2):220–231. doi:10.1016/j.devcel.2010.07.007

6. Vastenhouw NL, Zhang Y, Woods IG, et al. Chromatin signature of embryonic pluripotency is established during genome activation. Nature. 2010;464(7290):922–926. doi:10.1038/nature08866

7. Zhang Y, Vastenhouw NL, Feng J, et al. Canonical nucleosome organization at promoters forms during genome activation. Genome Res. 2014;24(2):260–266. doi:10.1101/gr.157750.113

8. Dahl JA, Jung I, Aanes H, et al. Broad histone H3K4me3 domains in mouse oocytes modulate maternal-to-zygotic transition. Nature. 2016;537(7621):548–552. doi:10.1038/nature19360

9. Zhang B, Zheng H, Huang B, et al. Allelic reprogramming of the histone modification H3K4me3 in early mammalian development. Nature. 2016;537(7621):553–557. doi:10.1038/nature19361

10. Xia W, Xu J, Yu G, et al. Resetting histone modifications during human parental-to-zygotic transition. Science. 2019;365(6451):353–360. doi:10.1126/science.aaw5118

11. Laranjeira AC, Berger S, Kohlbrenner T, Greter NR, Hajnal A. Nutritional vitamin B12 regulates RAS/MAPK-mediated cell fate decisions through one-carbon metabolism. Nat Commun. 2024;15(1):8178. doi:10.1038/s41467-024-52556-3

12. Xiao Y, Bedet C, Robert VJP, et al. Caenorhabditis elegans chromatin-associated proteins SET-2 and ASH-2 are differentially required for histone H3 Lys 4 methylation in embryos and adult germ cells. Proceedings of the National Academy of Sciences. 2011;108(20):8305–8310. doi:10.1073/pnas.1019290108

13. Käser-Pébernard S, Müller F, Wicky C. LET-418/Mi2 and SPR-5/LSD1 Cooperatively Prevent Somatic Reprogramming of C. elegans Germline Stem Cells. Stem Cell Reports. 2014;2(4):547–559. doi:10.1016/j.stemcr.2014.02.007

14. Dubois ML, Boisvert FM. The Nucleolus: Structure and Function. In: Bazett-Jones DP, Dellaire G, eds. The Functional Nucleus. Springer International Publishing; 2016:29–49. doi:10.1007/978-3-319-38882-3_2

15. Korčeková D, Gombitová A, Raška I, Cmarko D, Lanctôt C. Nucleologenesis in the Caenorhabditis elegans Embryo. PLOS ONE. 2012;7(7):e40290. doi:10.1371/journal.pone.0040290

16. Bjerregaard B, Maddox-Hyttel P. Regulation of ribosomal RNA gene expression in porcine oocytes. Animal Reproduction Science. 2004;82-83:605–616. doi:10.1016/j.anireprosci.2004.04.023

17. Fuchs J, Loidl J. Behaviour of nucleolus organizing regions (NORs) and nucleoli during mitotic and meiotic divisions in budding yeast. Chromosome Res. 2004;12(5):427–438. doi:10.1023/B:CHRO.0000034726.05374.db

18. Fulka H, Rychtarova J, Loi P. The nucleolus-like and precursor bodies of mammalian oocytes and embryos and their possible role in post-fertilization centromere remodelling. Biochemical Society Transactions. 2020;48(2):581–593. doi:10.1042/BST20190847

19. Kresoja-Rakic J, Santoro R. Nucleolus and rRNA Gene Chromatin in Early Embryo Development. Trends in Genetics. 2019;35(11):868–879. doi:10.1016/j.tig.2019.06.005

20. Zatsepina OV, Bouniol-Baly C, Amirand C, Debey P. Functional and Molecular Reorganization of the Nucleolar Apparatus in Maturing Mouse Oocytes. Developmental Biology. 2000;223(2):354–370. doi:10.1006/dbio.2000.9762

21. Xu D, Chen X, Kuang Y, et al. rRNA intermediates coordinate the formation of nucleolar vacuoles in C. elegans. Cell Reports. 2023;42(8). doi:10.1016/j.celrep.2023.112915

22. Cenik ES, Meng X, Tang NH, et al. Maternal Ribosomes Are Sufficient for Tissue Diversification during Embryonic Development in C. elegans. Developmental Cell. 2019;48(6):811–826.e6. doi:10.1016/j.devcel.2019.01.019

23. Sims J, Copenhaver GP, Schlögelhofer P. Meiotic DNA Repair in the Nucleolus Employs a Nonhomologous End-Joining Mechanism. The Plant Cell. 2019;31(9):2259–2275. doi:10.1105/tpc.19.00367

24. Fulka H, Langerova A. The maternal nucleolus plays a key role in centromere satellite maintenance during the oocyte to embryo transition. Development. 2014;141(8):1694–1704. doi:10.1242/dev.105940

25. Belew MD, Chien E, Michael WM. Characterization of factors that underlie transcriptional silencing in C. elegans oocytes. PLOS Genetics. 2023;19(7):e1010831. doi:10.1371/journal.pgen.1010831

26. Turowski TW, Tollervey D. Cotranscriptional events in eukaryotic ribosome synthesis. WIREs RNA. 2015;6(1):129–139. doi:10.1002/wrna.1263

27. Dash S, Lamb MC, Lange JJ, et al. rRNA transcription is integral to phase separation and maintenance of nucleolar structure. PLOS Genetics. 2023;19(8):e1010854. doi:10.1371/journal.pgen.1010854

28. Lafontaine DLJ, Riback JA, Bascetin R, Brangwynne CP. The nucleolus as a multiphase liquid condensate. Nat Rev Mol Cell Biol. 2021;22(3):165–182. doi:10.1038/s41580-020-0272-6

29. Mukherjee S, Firpo EJ, Wang Y, Roberts JM. Separation of telomerase functions by reverse genetics. Proceedings of the National Academy of Sciences. 2011;108(50):E1363–E1371. doi:10.1073/pnas.1112414108

30. Son MY, Belan O, Spirek M, et al. RAD51 separation of function mutation disables replication fork maintenance but preserves DSB repair. iScience. 2024;27(4). doi:10.1016/j.isci.2024.109524

31. Taddei A, Hediger F, Neumann FR, Bauer C, Gasser SM. Separation of silencing from perinuclear anchoring functions in yeast Ku80, Sir4 and Esc1 proteins. The EMBO Journal. 2004;23(6):1301–1312. doi:10.1038/sj.emboj.7600144

32. Ginisty H, Amalric F, Bouvet P. Nucleolin functions in the first step of ribosomal RNA processing. The EMBO Journal. 1998;17(5):1476–1486. doi:10.1093/emboj/17.5.1476

33. Roger B, Moisand A, Amalric F, Bouvet P. Nucleolin provides a link between RNA polymerase I transcription and pre-ribosome assembly. Chromosoma. 2003;111(6):399–407. doi:10.1007/s00412-002-0221-5

34. Spaulding EL, Feidler AM, Cook LA, Updike DL. RG/RGG repeats in the C. elegans homologs of Nucleolin and GAR1 contribute to sub-nucleolar phase separation. Nat Commun. 2022;13(1):6585. doi:10.1038/s41467-022-34225-5

35. Lee CC, Tsai YT, Kao CW, et al. Mutation of a Nopp140 gene dao-5 alters rDNA transcription and increases germ cell apoptosis in C. elegans. Cell Death Dis. 2014;5(4):e1158–e1158. doi:10.1038/cddis.2014.114

36. Rog O, Dernburg AF. Direct Visualization Reveals Kinetics of Meiotic Chromosome Synapsis. Cell Reports. 2015;10(10):1639–1645. doi:10.1016/j.celrep.2015.02.032

37. Smolikov S, Eizinger A, Schild-Prufert K, et al. SYP-3 Restricts Synaptonemal Complex Assembly to Bridge Paired Chromosome Axes During Meiosis in Caenorhabditis elegans. Genetics. 2007;176(4):2015–2025. doi:10.1534/genetics.107.072413

38. Eberhard R, Stergiou L, Hofmann ER, et al. Ribosome Synthesis and MAPK Activity Modulate Ionizing Radiation-Induced Germ Cell Apoptosis in Caenorhabditis elegans. PLOS Genetics. 2013;9(11):e1003943. doi:10.1371/journal.pgen.1003943

39. McCarter J, Bartlett B, Dang T, Schedl T. On the Control of Oocyte Meiotic Maturation and Ovulation in*Caenorhabditis elegans*. Developmental Biology. 1999;205(1):111–128. doi:10.1006/dbio.1998.9109

40. Berry J, Weber SC, Vaidya N, Haataja M, Brangwynne CP. RNA transcription modulates phase transition-driven nuclear body assembly. Proceedings of the National Academy of Sciences. 2015;112(38):E5237–E5245. doi:10.1073/pnas.1509317112

41. Liao S, Chen X, Xu T, et al. Antisense ribosomal siRNAs inhibit RNA polymerase I-directed transcription in C. elegans. Nucleic Acids Research. 2021;49(16):9194–9210. doi:10.1093/nar/gkab662

42. Korsholm LM, Gál Z, Nieto B, et al. Recent advances in the nucleolar responses to DNA double-strand breaks. Nucleic Acids Research. 2020;48(17):9449–9461. doi:10.1093/nar/gkaa713

43. Zhao Q, Rangan R, Weng S, Özdemir C, Cenik ES. Inhibition of ribosome biogenesis in the epidermis is sufficient to trigger organism-wide growth quiescence independently of nutritional status in C. elegans. PLOS Biology. 2023;21(8):e3002276. doi:10.1371/journal.pbio.3002276

44. Nousch M, Eckmann CR. Translational Control in the Caenorhabditis elegans Germ Line. In: Schedl T, ed. Germ Cell Development in C. Elegans. Springer; 2013:205–247. doi:10.1007/978-1-4614-4015-4_8

45. Racher H, Hansen D. Translational control in the C. elegans hermaphrodite germ line. Genome. 2010;53(2):83–102. doi:10.1139/G09-090

46. Pushpa K, Kumar GA, Subramaniam K. Translational Control of Germ Cell Decisions. Results Probl Cell Differ. 2017;59:175–200. doi:10.1007/978-3-319-44820-6_6

47. Iouk TL, Aitchison JD, Maguire S, Wozniak RW. Rrb1p, a Yeast Nuclear WD-Repeat Protein Involved in the Regulation of Ribosome Biosynthesis. Molecular and Cellular Biology. 2001;21(4):1260–1271. doi:10.1128/MCB.21.4.1260-1271.2001

48. Schütz S, Fischer U, Altvater M, et al. A RanGTP-independent mechanism allows ribosomal protein nuclear import for ribosome assembly. Hegde RS, ed. eLife. 2014;3:e03473. doi:10.7554/eLife.03473

49. Awad D, Prattes M, Kofler L, et al. Inhibiting eukaryotic ribosome biogenesis. BMC Biology. 2019;17(1):46. doi:10.1186/s12915-019-0664-2

50. Bass HW, Marshall WF, Sedat JW, Agard DA, Cande WZ. Telomeres Cluster De Novo before the Initiation of Synapsis: A Three-dimensional Spatial Analysis of Telomere Positions before and during Meiotic Prophase. Journal of Cell Biology. 1997;137(1):5–18. doi:10.1083/jcb.137.1.5

51. Chriss A, Börner GV, Ryan SD. Agent-based modeling of nuclear chromosome ensembles identifies determinants of homolog pairing during meiosis. PLOS Computational Biology. 2024;20(5):e1011416. doi:10.1371/journal.pcbi.1011416

52. MacQueen AJ, Villeneuve AM. Nuclear reorganization and homologous chromosome pairing during meiotic prophase require *C. elegans chk-2*. Genes Dev. 2001;15(13):1674–1687. doi:10.1101/gad.902601

53. Reinke V, Gil IS, Ward S, Kazmer K. Genome-wide germline-enriched and sex-biased expression profiles in Caenorhabditis elegans. Development. 2004;131(2):311–323. doi:10.1242/dev.00914

54. Chiu TP, Xin B, Markarian N, Wang Y, Rohs R. TFBSshape: an expanded motif database for DNA shape features of transcription factor binding sites. Nucleic Acids Res. 2020;48(D1):D246–D255. doi:10.1093/nar/gkz970

55. Chi W, Reinke V. Promotion of oogenesis and embryogenesis in the C. elegans gonad by EFL-1/DPL-1 (E2F) does not require LIN-35 (pRB). Development. 2006;133(16):3147–3157. doi:10.1242/dev.02490

56. Angus-Hill ML, Schlichter A, Roberts D, Erdjument-Bromage H, Tempst P, Cairns BR. A Rsc3/Rsc30 Zinc Cluster Dimer Reveals Novel Roles for the Chromatin Remodeler RSC in Gene Expression and Cell Cycle Control. Molecular Cell. 2001;7(4):741–751. doi:10.1016/S1097-2765(01)00219-2

57. Albert B, Knight B, Merwin J, et al. A Molecular Titration System Coordinates Ribosomal Protein Gene Transcription with Ribosomal RNA Synthesis. Molecular Cell. 2016;64(4):720–733. doi:10.1016/j.molcel.2016.10.003

58. Han M, Wei G, McManus CE, Hillier LW, Reinke V. Isolated C. elegans germ nuclei exhibit distinct genomic profiles of histone modification and gene expression. BMC Genomics. 2019;20:500. doi:10.1186/s12864-019-5893-9

59. McManus CE, Mazzetto M, Wei G, Han M, Reinke V. The zinc-finger protein OEF-1 stabilizes histone modification patterns and promotes efficient splicing in the Caenorhabditis elegans germline. G3 (Bethesda). 2021;11(12):jkab329. doi:10.1093/g3journal/jkab329

60. Tabuchi TM, Rechtsteiner A, Jeffers TE, Egelhofer TA, Murphy CT, Strome S. Caenorhabditis elegans sperm carry a histone-based epigenetic memory of both spermatogenesis and oogenesis. Nat Commun. 2018;9(1):4310. doi:10.1038/s41467-018-06236-8

61. Li T, Kelly WG. A Role for Set1/MLL-Related Components in Epigenetic Regulation of the Caenorhabditis elegans Germ Line. Schübeler D, ed. PLoS Genet. 2011;7(3):e1001349. doi:10.1371/journal.pgen.1001349

62. Nottke AC, Beese-Sims SE, Pantalena LF, Reinke V, Shi Y, Colaiácovo MP. SPR-5 is a histone H3K4 demethylase with a role in meiotic double-strand break repair. Proc Natl Acad Sci USA. 2011;108(31):12805–12810. doi:10.1073/pnas.1102298108

63. Herbette M, Mercier MG, Michal F, et al. The *C. elegans* SET-2/SET1 histone H3 Lys4 (H3K4) methyltransferase preserves genome stability in the germline. DNA Repair. 2017;57:139–150. doi:10.1016/j.dnarep.2017.07.007

64. Herbette M, Robert V, Bailly A, et al. A Role for Caenorhabditis elegans COMPASS in Germline Chromatin Organization. Cells. 2020;9(9):2049. doi:10.3390/cells9092049

65. Freeman TF, Zhao Q, Surya A, Rothe R, Cenik ES. Ribosome biogenesis disruption mediated chromatin structure changes revealed by SRAtac, a customizable end to end analysis pipeline for ATAC-seq. BMC Genomics. 2023;24(1):512. doi:10.1186/s12864-023-09576-y

66. Al-Refaie N, Padovani F, Hornung J, et al. Fasting shapes chromatin architecture through an mTOR/RNA Pol I axis. Nat Cell Biol. 2024;26(11):1903–1917. doi:10.1038/s41556-024-01512-w

67. Bersaglieri C, Kresoja-Rakic J, Gupta S, et al. Genome-wide maps of nucleolus interactions reveal distinct layers of repressive chromatin domains. Nat Commun. 2022;13(1):1483. doi:10.1038/s41467-022-29146-2

68. Hua L, Yan D, Wan C, Hu B. Nucleolus and Nucleolar Stress: From Cell Fate Decision to Disease Development. Cells. 2022;11(19):3017. doi:10.3390/cells11193017

69. Paredes S, Maggert KA. Ribosomal DNA contributes to global chromatin regulation. Proceedings of the National Academy of Sciences. 2009;106(42):17829–17834. doi:10.1073/pnas.0906811106

70. Wang C, Ma H, Baserga SJ, Pederson T, Huang S. Nucleolar structure connects with global nuclear organization. MBoC. 2023;34(12):ar114. doi:10.1091/mbc.E23-02-0062

71. Potapova TA, Gerton JL. Ribosomal DNA and the nucleolus in the context of genome organization. Chromosome Res. 2019;27(1):109–127. doi:10.1007/s10577-018-9600-5

72. Bagijn MP, Goldstein LD, Sapetschnig A, et al. Function, Targets, and Evolution of Caenorhabditis elegans piRNAs. Science. 2012;337(6094):574–578. doi:10.1126/science.1220952

73. Gu W, Shirayama M, Conte D, et al. Distinct Argonaute-Mediated 22G-RNA Pathways Direct Genome Surveillance in the *C. elegans* Germline. Molecular Cell. 2009;36(2):231–244. doi:10.1016/j.molcel.2009.09.020

74. Shen EZ, Chen H, Ozturk AR, et al. Identification of piRNA Binding Sites Reveals the Argonaute Regulatory Landscape of the *C. elegans* Germline. Cell. 2018;172(5):937–951.e18. doi:10.1016/j.cell.2018.02.002

75. Belew MD, Chien E, Wong M, Michael WM. The topoisomerase II/condensin II axis silences transcription during germline specification in *Caenorhabditis elegans*. Arbeitman M, ed. G3: Genes, Genomes, Genetics. Published online October 3, 2024:jkae236. doi:10.1093/g3journal/jkae236

76. Crittenden SL, Bernstein DS, Bachorik JL, et al. A conserved RNA-binding protein controls germline stem cells in Caenorhabditis elegans. Nature. 2002;417(6889):660–663. doi:10.1038/nature754

77. Diag A, Schilling M, Klironomos F, Ayoub S, Rajewsky N. Spatiotemporal m(i)RNA Architecture and 3′ UTR Regulation in the *C. elegans* Germline. Developmental Cell. 2018;47(6):785–800.e8. doi:10.1016/j.devcel.2018.10.005

78. Merritt C, Rasoloson D, Ko D, Seydoux G. 3′ UTRs Are the Primary Regulators of Gene Expression in the C. elegans Germline. Current Biology. 2008;18(19):1476–1482. doi:10.1016/j.cub.2008.08.013

79. Wickens M, Bernstein DS, Kimble J, Parker R. A PUF family portrait: 3′UTR regulation as a way of life. Trends in Genetics. 2002;18(3):150–157. doi:10.1016/S0168-9525(01)02616-6

80. Deisenroth C, Franklin DA, Zhang Y. The Evolution of the Ribosomal Protein–MDM2–p53 Pathway. Cold Spring Harb Perspect Med. 2016;6(12):a026138. doi:10.1101/cshperspect.a026138

81. Hannan KM, Soo P, Wong MS, et al. Nuclear stabilization of p53 requires a functional nucleolar surveillance pathway. Cell Reports. 2022;41(5):111571. doi:10.1016/j.celrep.2022.111571

82. Ma H, Pederson T. The nucleolus stress response is coupled to an ATR-Chk1–mediated G2 arrest. MBoC. 2013;24(9):1334–1342. doi:10.1091/mbc.e12-12-0881

83. Li W, Yanowitz JL. ATM and ATR Influence Meiotic Crossover Formation Through Antagonistic and Overlapping Functions in Caenorhabditis elegans. Genetics. 2019;212(2):431–443. doi:10.1534/genetics.119.302193

84. Yu Z, Kim HJ, Dernburg AF. ATM signaling modulates cohesin behavior in meiotic prophase and proliferating cells. Nat Struct Mol Biol. 2023;30(4):436–450. doi:10.1038/s41594-023-00929-5

85. Santoro R, Grummt I. Molecular Mechanisms Mediating Methylation-Dependent Silencing of Ribosomal Gene Transcription. Molecular Cell. 2001;8(3):719–725. doi:10.1016/S1097-2765(01)00317-3

86. Drygin D, Lin A, Bliesath J, et al. Targeting RNA Polymerase I with an Oral Small Molecule CX-5461 Inhibits Ribosomal RNA Synthesis and Solid Tumor Growth. Cancer Res. 2011;71(4):1418–1430. doi:10.1158/0008-5472.CAN-10-1728

87. Nechay M, Wang D, Kleiner RE. Inhibition of nucleolar transcription by oxaliplatin involves ATM/ATR kinase signaling. Cell Chemical Biology. 2023;30(8):906–919.e4. doi:10.1016/j.chembiol.2023.06.010

88. Xu H, Di Antonio M, McKinney S, et al. CX-5461 is a DNA G-quadruplex stabilizer with selective lethality in BRCA1/2 deficient tumours. Nat Commun. 2017;8(1):14432. doi:10.1038/ncomms14432

89. Osmanoğulları SC, Forough M, Persil Çetinkol Ö, Arslan Udum Y, Toppare L. Electrochemical detection of Oxaliplatin induced DNA damage in G-quadruplex structures. Analytical Biochemistry. 2023;671:115149. doi:10.1016/j.ab.2023.115149

90. Fulka H, Mrazek M, Fulka J. Nucleolar dysfunction may be associated with infertility in humans. Fertility and Sterility. 2004;82(2):486–487. doi:10.1016/j.fertnstert.2003.12.042

91. Sanchez E, Laplace-Builhé B, Mau-Them FT, et al. POLR1B and neural crest cell anomalies in Treacher Collins syndrome type 4. Genet Med. 2020;22(3):547–556. doi:10.1038/s41436-019-0669-9

92. Trainor PA. Craniofacial birth defects: The role of neural crest cells in the etiology and pathogenesis of Treacher Collins syndrome and the potential for prevention. American Journal of Medical Genetics Part A. 2010;152A(12):2984–2994. doi:10.1002/ajmg.a.33454

93. Dickinson DJ, Pani AM, Heppert JK, Higgins CD, Goldstein B. Streamlined Genome Engineering with a Self-Excising Drug Selection Cassette. Genetics. 2015;200(4):1035–1049. doi:10.1534/genetics.115.178335

94. R: A Language and Environment for Statistical Computing. Published online 2023. https://www.R-project.org/

95. Stirling DR, Swain-Bowden MJ, Lucas AM, Carpenter AE, Cimini BA, Goodman A. CellProfiler 4: improvements in speed, utility and usability. BMC Bioinformatics. 2021;22(1):433. doi:10.1186/s12859-021-04344-9

96. Schneider CA, Rasband WS, Eliceiri KW. NIH Image to ImageJ: 25 years of image analysis. Nat Methods. 2012;9(7):671–675. doi:10.1038/nmeth.2089

97. Martin M. Cutadapt removes adapter sequences from high-throughput sequencing reads. EMBnet.journal. 2011;17(1):10–12. doi:10.14806/ej.17.1.200

98. Kim D, Paggi JM, Park C, Bennett C, Salzberg SL. Graph-based genome alignment and genotyping with HISAT2 and HISAT-genotype. Nat Biotechnol. 2019;37(8):907–915. doi:10.1038/s41587-019-0201-4

99. Picard toolkit. Published online 2019. Accessed December 12, 2023. https://broadinstitute.github.io/picard/

100. Liao Y, Smyth GK, Shi W. featureCounts: an efficient general purpose program for assigning sequence reads to genomic features. Bioinformatics. 2014;30(7):923–930. doi:10.1093/bioinformatics/btt656

101. Ramírez F, Ryan DP, Grüning B, et al. deepTools2: a next generation web server for deep-sequencing data analysis. Nucleic Acids Research. 2016;44(W1):W160–W165. doi:10.1093/nar/gkw257

102. Hahne F, Durinck S, Ivanek R, et al. Gviz: Plotting data and annotation information along genomic coordinates. Published online 2023. doi:10.18129/B9.bioc.Gviz

103. Love MI, Huber W, Anders S. Moderated estimation of fold change and dispersion for RNA-seq data with DESeq2. Genome Biology. 2014;15(12):550. doi:10.1186/s13059-014-0550-8

104. Risso D, Ngai J, Speed TP, Dudoit S. Normalization of RNA-seq data using factor analysis of control genes or samples. Nat Biotechnol. 2014;32(9):896–902. doi:10.1038/nbt.2931

105. Chen Y, Lun ATL, Smyth GK. From reads to genes to pathways: differential expression analysis of RNA-Seq experiments using Rsubread and the edgeR quasi-likelihood pipeline. Published online August 2, 2016. doi:10.12688/f1000research.8987.2

106. Zhu A, Ibrahim JG, Love MI. Heavy-tailed prior distributions for sequence count data: removing the noise and preserving large differences. Bioinformatics. 2019;35(12):2084–2092. doi:10.1093/bioinformatics/bty895

107. Wang Q, Li M, Wu T, et al. Exploring Epigenomic Datasets by ChIPseeker. Current Protocols. 2022;2(10):e585. doi:10.1002/cpz1.585

108. Annotation package for TxDb object(s). Published online 2019. Accessed December 12, 2023. http://bioconductor.org/packages/TxDb.Celegans.UCSC.ce11.refGene/

109. Wu T, Hu E, Xu S, et al. clusterProfiler 4.0: A universal enrichment tool for interpreting omics data. The Innovation. 2021;2(3):100141. doi:10.1016/j.xinn.2021.100141

110. Yu G, Wang LG, Han Y, He QY. clusterProfiler: an R Package for Comparing Biological Themes Among Gene Clusters. OMICS: A Journal of Integrative Biology. 2012;16(5):284–287. doi:10.1089/omi.2011.0118

111. Yu G, Gao CH. enrichplot: Visualization of Functional Enrichment Result. Published online 2024. https://yulab-smu.top/biomedical-knowledge-mining-book/

112. Bailey TL, Johnson J, Grant CE, Noble WS. The MEME Suite. Nucleic Acids Research. 2015;43(W1):W39–W49. doi:10.1093/nar/gkv416

113. Bailey TL. STREME: accurate and versatile sequence motif discovery. Bioinformatics. 2021;37(18):2834–2840. doi:10.1093/bioinformatics/btab203

114. Gupta S, Stamatoyannopoulos JA, Bailey TL, Noble WS. Quantifying similarity between motifs. Genome Biology. 2007;8(2):R24. doi:10.1186/gb-2007-8-2-r24

115. Lawrence M, Huber W, Pagès H, et al. Software for Computing and Annotating Genomic Ranges. PLOS Computational Biology. 2013;9(8):e1003118. doi:10.1371/journal.pcbi.1003118

116. SRA Toolkit Development Team. SRA Toolkit. https://trace.ncbi.nlm.nih.gov/Traces/sra/sra.cgi?view=software

117. Ewels PA, Peltzer A, Fillinger S, et al. The nf-core framework for community-curated bioinformatics pipelines. Nat Biotechnol. 2020;38(3):276–278. doi:10.1038/s41587-020-0439-x

118. nf-core/chipseq. 10.5281/zenodo.3240506

119. Kent J, UCSC Genome Browser Group. faCount. https://hgdownload.soe.ucsc.edu/admin/exe/linux.x86_64/faCount

120. Bankhead P, Loughrey MB, Fernández JA, et al. QuPath: Open source software for digital pathology image analysis. Sci Rep. 2017;7(1):16878. doi:10.1038/s41598-017-17204-5

121. Schmidt U, Weigert M, Broaddus C, Myers G. Cell Detection with Star-Convex Polygons. In: Frangi AF, Schnabel JA, Davatzikos C, Alberola-López C, Fichtinger G, eds. Medical Image Computing and Computer Assisted Intervention – MICCAI 2018. Springer International Publishing; 2018:265–273. doi:10.1007/978-3-030-00934-2_30

122. Toraason E, Adler VL, Kurhanewicz NA, et al. Automated and customizable quantitative image analysis of whole Caenorhabditis elegans germlines. Genetics. 2021;217(3):iyab010. doi:10.1093/genetics/iyab010

